# Fibroblasts neurotrophin signaling sustains pathological vascular maturation in rheumatoid arthritis

**DOI:** 10.64898/2026.03.12.711120

**Authors:** Vikram Khedgikar, Qian Qin, Miles Tran, Shideh Kazerounian, Gao Ce, Sean A Prell, Alexa B. R. McIntyre, Kartik Bhamidipati, Sonia Presti, Philip E Blazar, Jeffrey K Lange, Morgan H Jones, Ellen M Gravallese, Mihir D Wechalekar, Kevin Wei

## Abstract

Treatment failures in rheumatoid arthritis (RA) leads to undesirable morbidity associated with immunosuppression. Recent studies of synovial tissue from refractory RA patients highlight the role of synovial fibroblasts and vascular endothelium in driving treatment failure. Utilizing high-dimensional spatial transcriptomics, we uncovered a crucial role for neurotrophin signaling in driving abnormal vascular maturation in RA synovia. Neurotrophins, including nerve growth factor (NGF), brain-derived neurotrophic factor (BDNF), and neurotrophin-3 (NT3), induce differentiation of synovial fibroblasts into mural cells - pericytes and vascular smooth muscle cells. Mechanistically, NOTCH3 signaling activates a cascade of neurotrophin signaling through transcriptional induction of NGFR, a co-receptor for NGF. In RA synovial tissue explants, stimulation with NGF, BDNF, or NT3 leads to a dramatic increase in maturation of synovial tissue vasculature. Conversely, pharmacologic inhibition with neurotrophin inhibitors drastically abolished maturation of vascularization in RA synovial explants. Notably, the FDA-approved tropomyosin receptor kinase (TRK) inhibitors larotrectinib and entrectinib effectively reverse synovial vascular maturation in human RA tissue explants.Our findings suggest that fibroblast-derived neurotrophin signaling is a critical pathway in sustaining mature blood vessels in RA synovia, and that neurotrophin inhibitors reverse abnormal vascular maturation in RA.

**One Sentence Summary:** In rheumatoid arthritis, fibroblast neurotrophin signaling drives abnormal vascular maturation by inducing differentiation of fibroblasts into vascular mural cells.

## Introduction

Rheumatoid arthritis (RA) is a chronic inflammatory disease characterized by hyperplasia of synovial tissue, leading to irreversible joint damage. Treatment failures in RA lead to high costs to society and cause undesirable morbidity associated with immunosuppression. Our previous study utilizing high-dimensional analysis of pre- and post-treatment RA synovial biopsies suggest that while immunosuppression depletes infiltrating immune cells in RA synovia, the stromal component undergoes pathological remodeling (*1*). A key unmet need and fundamental question remains on cellular and molecular pathways that sustain pathological stromal remodeling in RA.

The blood vessel is composed of endothelial and mural cells -pericytes and vascular smooth muscle cells (VSMCs), and vascular fibroblasts(*2*),(*3*),(*4*). Recruitment of mural cells to endothelium is considered the final step of vascular maturation where the nascent endothelium becomes functional blood vessels(*5*). Mural cells play critical roles in maintaining the integrity and function of blood vessels(*6*),(*7*),(*8*),(*9*). Disruption of mural cell homeostasis is a central pathological feature underlying vascular diseases such as pulmonary arterial hypertension(*10*) atherosclerosis(*3*),(*11*), and age-related small-vessel disease of the central nervous system(*12*). In RA synovia, the vasculature undergoes dramatic remodeling in response to chronic inflammation and hypoxia, resulting in formation of new blood vessels and maturation of newly formed vessels(*13*). In addition to their role in supporting the pathological growth of synovia, vascular endothelial cells play an important role in driving pathogenic fibroblast expansion(*14*) and axonal sprouting(*15*) in RA. An important unanswered question in RA is what pathway sustains pathological vascular remodeling in RA, and whether current immunosuppressive therapies are sufficient to reverse pathological vascularization in RA. In our previous study(*16*), we demonstrated a role for NOTCH3 in vascular fibroblast differentiation and that fibrogenic signals from synovial endothelial cells contribute to treatment-resistance in RA (*1, 14*). Here, utilizing high-dimensional spatial transcriptomics to profile RA synovial microvasculature and uncovered an unexpected role for neurotrophin receptor signaling in pathological vascular maturation in RA synovia.

## Results

### Pathological synovial vascular maturation in RA

To comprehensively characterize the microvasculature in RA synovia before and after immunosuppressive treatment, we expanded our spatial transcriptomic analysis from a cohort of treatment-naive patients(*1*) which includes paired biopsies from pre-treatment and 6-month post- triple csDMARD therapy (Hydroxychloroquine, methotrexate, and sulfasalazine) or TNFinhibitor (TNFi:adalimumab) treatment (n = 22 RA patients, including longitudinal paired pre-treatment and 6-month post-treatment biopsies from all RA patients and 2 healthy donors). Following quality control and cell segmentation (see methods), we performed cell type annotation where each cell is mapped to the single-cell reference dataset generated from the AMP RA/SLE Consortium(*17*). At the cell lineage level, we identified 12 major synovial cell types, including B cells, T cells, NK, cells, myeloid cells, mast cells, proliferating cells, mural cells, endothelial cells, and synovial fibroblast subtypes (Fig. 1A and B). Next, we focused our analyses on 368,217 cells that make up the synovial microvasculature, including blood endothelial cells, lymphatic endothelial cells, pericytes, and vascular smooth muscle cells (VSMCs) (Fig. 1C). Based on expression of canonical endothelial cell and mural cell markers(*18*), we further classified cells into 6 fine-grain vascular cell types: (1) *POOXL*+arterial EC, (2) *VWF*+/*CO34*+venular EC, (3) *KOR*+ capillary EC, (4) *ACTA2*+ VSMCs, (5) *POGFRB*+ pericytes, and (6) *PROX1*+ lymphatic EC (Fig. 1C and D). Consistent with mature vascularization of synovial blood vessels in RA, spatial analysis confirmed pairing of endothelial cells with distinct mural cell subtypes, where arterial and capillary ECs are physically associated with VSMCs and pericytes, respectively (Fig. 1F).

**Figure 1.**
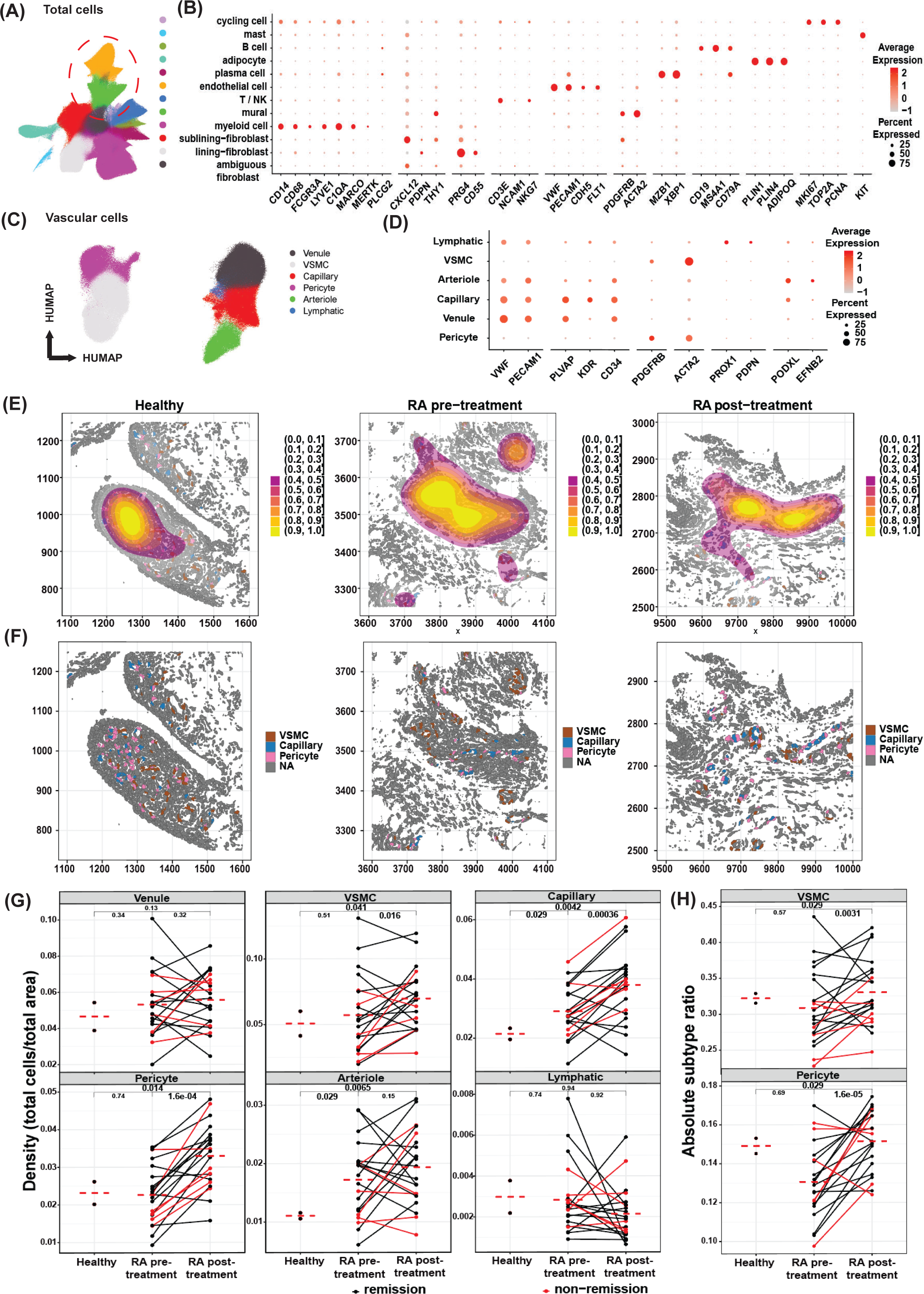
Persistent synovial vascular maturation in rheumatoid arthritis despite immunosuppressive therapy. High-dimensional spatial transcriptomic profiling was performed using the Xenium 5K Prime platform on 46 synovial tissue biopsy samples (22 RA patients and 2 healthy donors, including 22 RA patients with paired pre- and post-treatment biopsies). In total, 2,049,358 high-quality cells were analyzed, of which 368,217 cells were identified as vascular cells, comprising endothelial cells and mural cells. (A) UMAP projection of all cells, annotated with integrated cell-type labels. (B) Heatmap showing expression of marker genes used to define the identified cell clusters, dot color represents the average expression level, dot size is the expressed cell proportion, (C) UMAP projection of vascular cells only, annotated with integrated vascular subtype labels. (D) Heatmap showing marker gene expression for the identified vascular cell clusters. (E) Representative spatial plots of synovial tissue biopsies from healthy donors and RA patients before and after treatment, colored by cell type and illustrating vascular density. (F) Representative spatial plot highlighting vascular cells within synovial tissue. (G) Quantification of changes in overall vascular cell density defined as cell number per area across healthy, pre-treatment RA, and post-treatment RA samples. (H) Quantification of absolute cell proportion changes in vascular smooth muscle cells (VSMCs) and pericytes across healthy, pre-treatment, and post-treatment RA samples. Statistical analysis was performed using the Wilcoxon matched-pairs signed-rank test for paired patient samples.

To characterize pathological changes in RA synovial microvasculature, we first quantified the global microvascular density for each patient biopsy where the microvascular density is calculated by enumerating proportion of vascular cells as a function of total surface area of each synovial biopsy (Fig. 1E and F). Next, we quantified the microvascular density of each vascular cell subtype and then compared changes in microvascular density between healthy, pre-treatment RA, and post-treatment RA (Fig. 1G). When compared to healthy synovia, pre-treatment RA synovia is characterized by expansion of capillaries *(p = 0.029*) and arterioles *(p = 0.029)* (Fig. 1G), consistent with neovascularization and angiogenesis in the setting of synovial hyperplasia in RA. Surprisingly, we found that 6 months of immunosuppressive treatment with either triple csDMARD therapy or TNFi did not reduce the density of synovial microvasculature. Instead, we observed statistically significant interval increase in density of capillary ECs (*p = 0.00036; and p=0.004*2 compared to healthy), arteriolar ECs (*p = 0.0065*, compared to healthy), pericytes (*p=1.6e-05, p = 0.029* compared to healthy), and VSMCs (*p = 0.0031, p=0.029* compared to healthy) after 6 months of treatment (Fig. 1G). To confirm the persistence of increased vascularization in post-treatment RA synovia, we further analyzed the abundance of each vascular cell type as a function of total synovial cells from each biopsy (Fig. 1H). Analysis of cellular composition confirmed significant expansion of VSMC (*p = 0.0031, p = 0.029* compared to healthy) and pericytes *(p = 1.6 e-05, p = 0.029* compared to healthy) after 6 months of treatment (Fig. 1H). Further, the increase in synovial vascular maturation is specific to the capillary and arterial endothelia as the densities of venular ECs and lymphatic ECs did not change significantly between pre- and post-treatment. Importantly, the interval increase in synovial microvascular density occurred in RA patients regardless of whether or not patients reached criteria for clinical remission, as defined by DAS28-ESR < 2.6, at 6 months after treatment. Together, our data suggest that pathological vascular maturation, characterized by expansion of capillary and arterioles with concomitant increase in mural cell coverage, persists in RA synovia despite immunosuppressive treatment with either triple DMARD therapy or TNFi.

### Identification of neurotrophin receptor signaling during vascular maturation in RA

The expansion of the mural cell compartment in RA suggests that pathological maturation synovial microvasculature persists despite immunosuppressive treatment. Since the recruitment of mural cells to endothelia is a key step in forming a functional blood vessel during vascular maturation(*5*), we focused our analysis on growth factor receptors expressed by mural cells in RA microvasculature in search of a targetable pathway in RA vascular maturation.

Unexpectedly, we detected expression of the neurotrophin receptors *NGFR* and *NTRK2* in synovial mural cells (Fig. 2A). Neurotrophins, including Nerve Growth Factor (NGF), Brain-Derived Neurotrophic Factor (BDNF), and Neurotrophin-3 (NT3), are essential for neuronal development and function(*19*). Neurotrophins bind to their receptors, tropomyosin receptor kinases (TRKs), and p75 neurotrophin receptors (p75NTR/NGFR)(*20*). In RA synovia, *NTRK2*, encoding TRKB, a receptor for BDNF, is expressed by VSMCs while *NGFR* is predominantly expressed by pericytes and *NOTCH3*+ vascular fibroblasts (Fig. 2A). To verify celltype-specific expression of neurotrophin receptors in synovial microvasculatuture, we developed and applied a high-sensitivity, custom spatial transcriptomic panel (Table. S, 1 to 3) covering genes encoding neurotrophin receptors and found that *NGFR* expression was preferentially expressed by pericytes, whereas the expression of *NTRK2* and *NTRK3* (encoding TRKC) were restricted to VSMCs (Fig. S1D, Fig. S1). *NTRK1*, encoding TRKA, were expressed, albeit at a very low levels, in pericytes and VSMCs (Fig. S2). Next, we validated the expression of neurotrophin receptors on synovial mural cells by RNA in-situ hybridization (RNAscope) (Fig. 2B) followed by immunohistochemistry (Fig. 2C). Together, these analyses revealed that neurotrophin receptors are expressed on mural cells in RA synovial microvasculature and their expression persists despite 6-months of immunosuppressive therapy.

**Figure 2.**
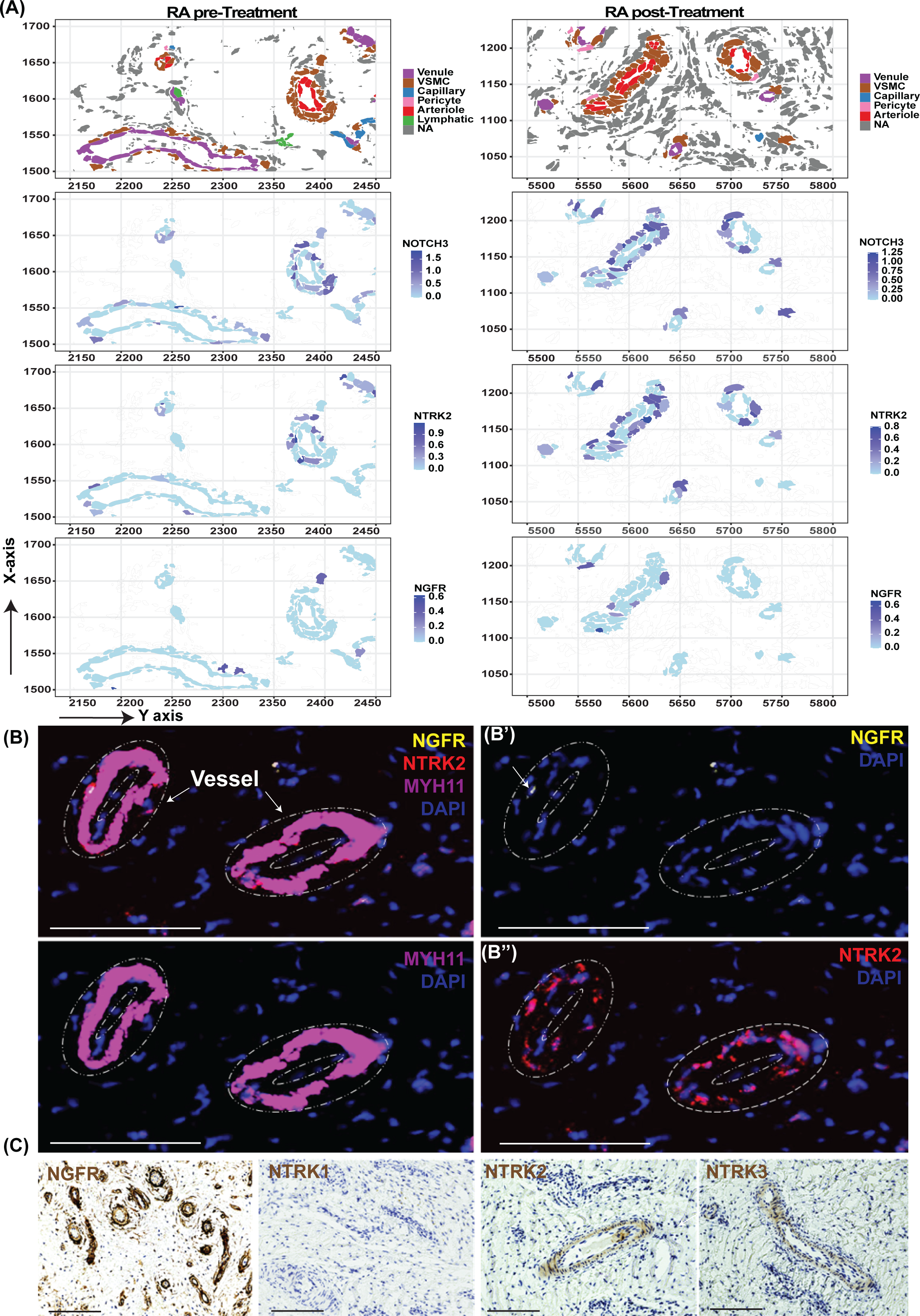
Mural cells in RA synovial microvasculature express neurotrophin receptors. (A) High-magnification spatial transcriptomic plots showing arterioles, venules, and capillaries in pre- and post-treatment RA synovial biopsy samples, colored by integrated vascular cell type annotations. The lower panels show high-magnification views of the same regions highlighting expression of NOTCH3 and neurotrophin receptors NGFR and NTRK2 in mural cells surrounding vascular structures. Relevant vascular regions are outlined.(B) RNAscope images showing transcript expression of the VSMC marker MYH11 (magenta) and neurotrophin receptors (B’) NGFR (yellow) and (B’’) NTRK2 (red) in RA synovial tissue sections, overlaid with DAPI (blue). Dotted lines indicate vascular structures defined by MYH11 expression.Scale bars, 150 µm. All images were acquired at 20x magnification and cropped to the same scale.(C) Immunohistochemistry images showing perivascular expression of neurotrophin receptors NGFR, NTRK1, NTRK2, and NTRK3 (brown) in RA synovial tissue. Scale bars, 150 µm. All images were acquired at 20x magnification and cropped to the same scale.

We have previously demonstrated that endothelial cells provide crucial signals in orchestrating synovial fibroblast differentiation in RA (2, 13). To test if endothelial cells induce neurotrophin signaling in synovial fibroblasts, we utilize a fibroblast-endothelial cell co-culture system where endothelial cells are mixed with synovial fibroblasts in a 1:3 ratio *in vitro* (Fig S3, Fig. 3A). In this context, fibroblasts in the proximal 1-2 cell layers away from the nearest endothelial cells expressed high levels of *NGFR, NTRK1, NTRK2, and NTRK3* (Fig. 3B). Consistent with induction of neurotrophin signaling, fibroblast-endothelial co-culture induced a significant increase in the production of neurotrophin ligands NGF (1.8 fold, *p = 0.002*)(Fig. S3B), and a trend towards increased production of BDNF and NT3. Consistent with our previous studies where *NOTCH3* plays a crucial role in endothelial-fibroblast crosstalk(*1, 14*), siRNA knockdown of NOTCH3 resulted in the downregulation of *NGFR, NTRK1, NTRK2*, and *NTRK3* in fibroblasts adjacent to endothelial cells (Fig. 3C, Fig. S7 A). Quantification of RNA transcripts showed siNOTCH3 significantly reduced expression of *NGFR* (55%, *p = 0.03*), *NTRK1* (47%, *p = 0.003*), *NTRK3* (62%, *p = 0.034)*, and a trend towards reduction in *NTRK2* (43%, *p = 0.053*) (Fig. S7, B to E). To test if the NOTCH pathway directly activates neurotrophin signaling in fibroblasts, we stimulated fibroblasts with NOTCH ligand Delta-like ligand 4 (DLL4). DLL4-stimulation upregulated the expression of VSMC marker *MYH11* (2.8-fold, *p=0.02*) and pericyte-marker *RGS5* (1.6-fold, *p = 0.005*). siRNA knockdown of *NTRK2* or *NTRK3* in fibroblasts ablated DLL4-induced *MYH11* expression (Fig. 3E), but had minimal effect on *RGS5* (Fig. S6, G and H). In contrast, knockdown of *NGFR* and *NTRK1* ablated DLL4-induced *RGS5* expression (Fig. 3D), but not *MYH11* (Fig S6, E and F). Together, these results indicate fibroblast neurotrophin signaling can be induced by endothelial cells via NOTCH3.

**Figure 3.**
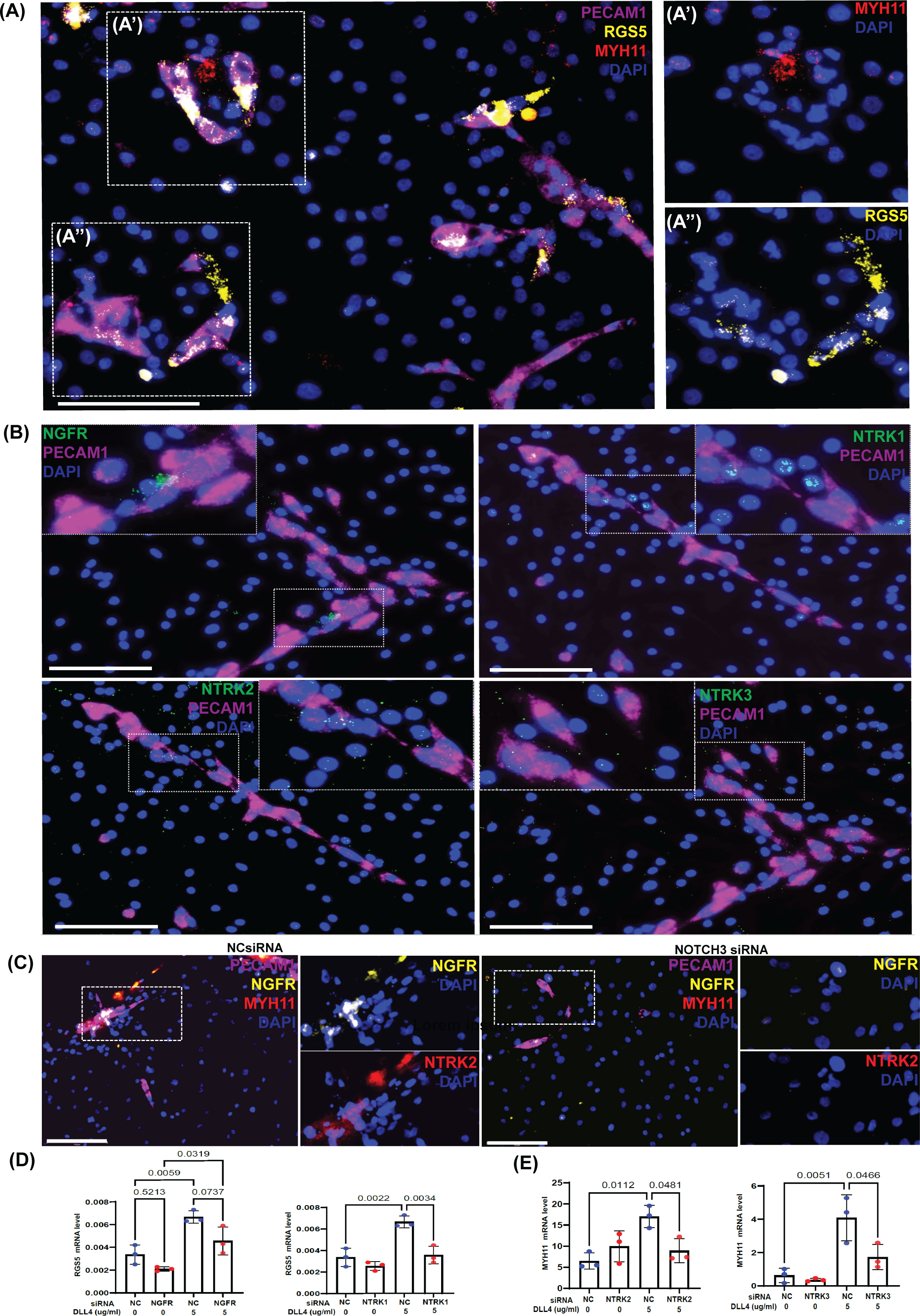
Endothelial cells induce expression of neurotrophin receptors in fibroblasts via NOTCH3 signaling. Human fibroblasts were co-cultured with human umbilical vein endothelial cells (HUVECs). Endothelial cells were identified by PECAM1 expression (magenta).(A) RNAscope images showing transcripts of VSMC marker MYH11(red) (A’) and the pericyte marker RGS5 (yellow) (A’’) and the in fibroblast-endothelial co-cultures, overlaid on DAPl(blue). Scale bar, 150 µm. All images were acquired at 20x magnification and cropped to the same scale. (B) RNAscope images showing transcripts of neurotrophin receptors NGFR, NTRK1, NTRK2, and NTRK3 (green) in the co-culture, overlaid on DAPl(blue). Enlarged cropped regions are shown in dotted boxes. Scale bar, 150 µm. All images were acquired at 20x magnification and cropped to the same scale.(C) Fibroblasts were treated with NOTCH3 siRNA and co-cultured with endothelial cells. RNAscope images show transcripts of NGFR (yellow) and NTRK2 (red) in the co-culture, overlaid on DAPl (blue). Enlarged cropped images are shown separately in dotted boxes. Scale bar, 150 µm. All images were acquired at 20x magnification and cropped to the same scale. (D-E) Fibroblasts were treated with siRNAs targeting NGFR, NTRK1, NTRK2, or NTRK3, in the presence or absence of DLL4 (5 µg/mL). (D) RGS5 mRNA expression following knockdown of NGFR or NTRK1. (E) MYH11 mRNA expression following knockdown of NTRK2 or NTRK3, in the presence or absence of DLL4. lndividual data points represent biological replicates; p values are indicated on the graphs. Statistical analysis was performed using a two tailed Student’s t test for comparisons between two groups and one-way ANOVA for comparisons among multiple groups. Post-hoc pairwise comparisons were conducted using t tests with Bonferroni correction to control for multiple comparisons.

### Neurotrophins induce differentiation of synovial fibroblasts into vascular mural cells

We next asked if neurotrophin signaling is sufficient to induce synovial fibroblasts to differentiate into vascular mural cells. Interestingly, neurotrophin stimulation exhibited shared and unique effects on fibroblast gene expression. Fibroblasts stimulated with NGF, BDNF, or NT3 exhibited marked induction of mural cell markers *ACTA2, SMTN, and TAGLN* (Fig. 4, A to C). NGF stimulation preferentially induced pericyte-specific markers *RGS5* (1.3-fold, *p = 0.0006*) and *ABCC9* (1.3-fold, *p = 0.002*). In contrast, NT3 stimulation induced VSMC-specific markers *CNN1* (1.2-fold, *p = 0.0001*) and *MYH11* (2.9-fold, *p = 0.0001*) (Fig. 4S A-B, Fig. 4C). BDNF stimulation induced both pericyte- and VSMC- markers (Fig. 4S A-B, Fig. 4B). Consistent with receptor specific activity of each neurotrophin, siRNA knockdown of *NGFR* or *NTRK1* reduced *RGS5* expression by approximately 50% (siNGFR, *p* = 0.002; siNTRK1, *p* = 0.0003) and completely abrogated NGF-induced *RGS5* expression (Fig. S4C). Conversely, knockdown of *NTRK2* or *NTRK3* abolished *MYH11* induction in response to BDNF and NT3 stimulation, respectively (Fig. S4D). Pharmacologic inhibition of TRKA (GW 441756)(*21, 22*) or TRKB (ANA-12)(*21*) similarly blocked NGF-induced and BDNF-induced *CNN1* expression (Fig. S4E and F).

**Figure 4.**
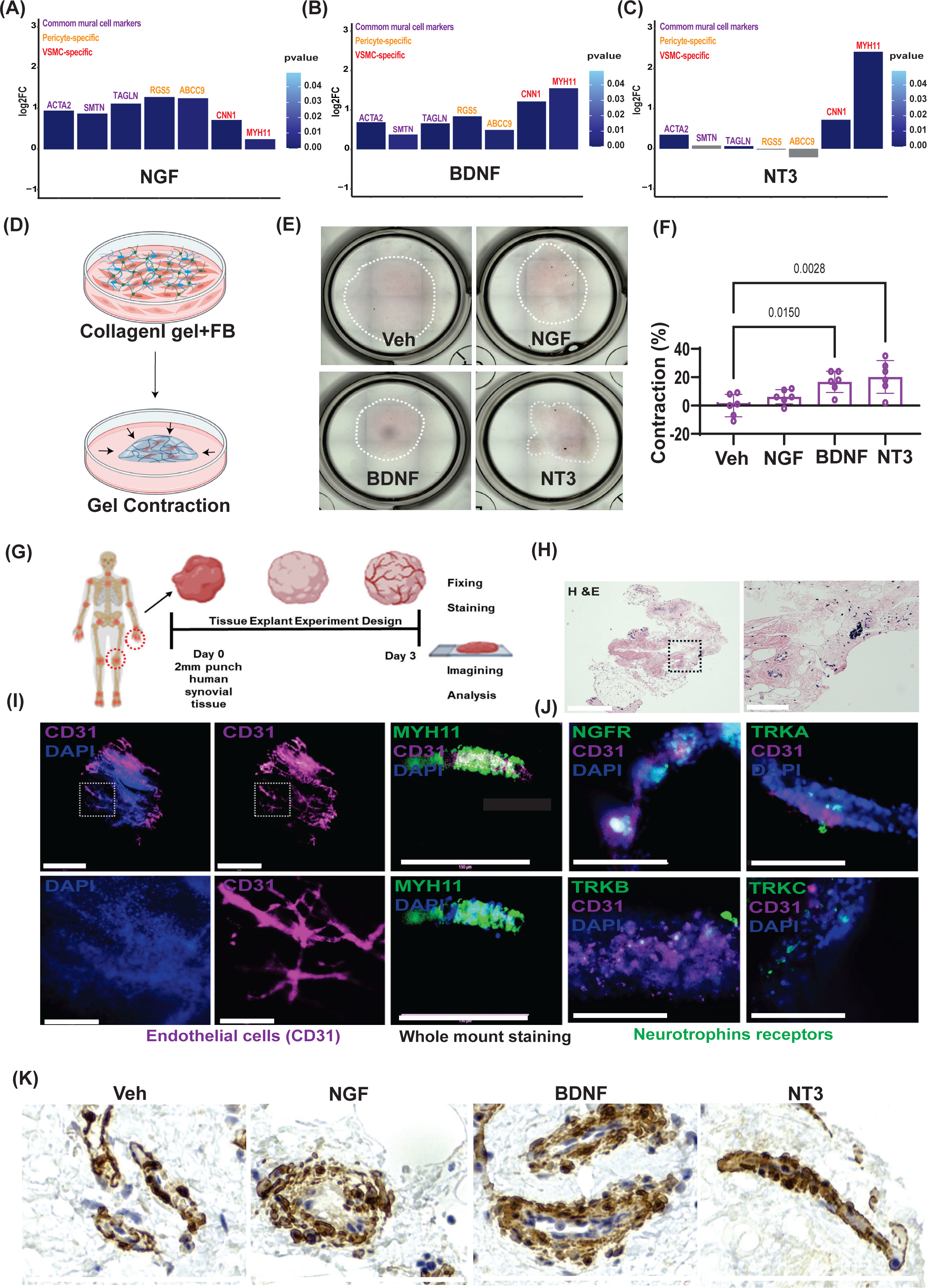
Neurotrophins induce fibroblast to mural cell differentiation. (A-C) Bar plots showing differential regulation of pericyte markers (RGS5, ABCC9), VSMC markers (CNN1, MYH11), and pan-mural cell markers (ACTA2, SMTN, TAGLN) in fibroblasts treated with (A) NGF, (B) BDNF, or (C) NT3 compared with untreated fibroblasts. Bar heights represent log₂ fold change, and bar color indicates the corresponding P values. (D) Schematic of the collagen gel contraction assay in which synovial fibroblasts were embedded in collagen I gels and treatment (E) Representative images of collagen I gel contraction following treatment with NGF, BDNF, or NT3 (F) Quantification of collagen I gel contraction expressed as percent contraction. Values represent mean ± standard deviation (SD). Individual data points indicate biological replicates. Statistical analysis was performed using one-way ANOVA, with P values shown. (G) Schematic illustrating the synovial organoid/explant experimental design. (H) Representative H&E image of synovial explant culture after 3 days of culture. (I) Whole-mount immunoluorescence images showing preservation of endothelial cells after 3 days in culture, marked by CD31 (magenta), and vascular smooth muscle cells marked by MYH11, surrounding CD31 vascular structures. (J) Whole-mount staining showing expression of neurotrophin receptors NGFR, NTRK1, NTRK2, and NTRK3 (green) around vascularature, together with CD31 (magenta) and DAPI (blue). (K) Synovial organoids embedded in Matrigel and treated with NGF, BDNF, or NT3, followed by fixation and staining for a-smooth muscle actin (aSMA) (brown) to visualize mural cells. All 20X images were cropped at similar scale. Scale bar, 150 µm.

To assess if fibroblast-derived mural cells acquire cellular functions specific to vascular mural cells, we first tested if fibroblast-derived mural cells could support vascular endothelial cell tube formation, a key function of mural cells(*23*). Stimulation with NGF, BDNF, or NT3 enhanced endothelial tubule formation compared to control. (Fig. S5A-B). We then tested if fibroblast-derived mural cells exhibit enhanced contractility, a key function of VSMCs(*24*)(*25*). Consistent with the ability of BDNF and NT3 to induce *MYH11* expression in fibroblasts, fibroblasts stimulated with BDNF or NT3 exhibited markedly enhanced contractility (Fig. S5B). We further validated the increased contractility using a collagen gel contraction assay(*12*). NT3 stimulation induced the greatest contractile response (21%, *p* = 0.002), followed by BDNF (16%, *p* = 0.01), whereas NGF had no significant effect on contractility (Fig. 4D-F, Fig S5C).

Given our observation that neurotrophins are able to induce synovial fibroblasts to differentiate into mural cells, we asked if fibroblasts in intact synovial tissue are capable of responding to neurotrophins and differentiate into mural cells *ex vivo*. To do this, we employed an *en-bloc* synovial tissue explant system(*1*) where synovial endothelial and mural cells retained lineage-specific marker expression (*PECAM1* and *MYH11*, respectively) and preserved tissue architecture for at least 3 days *ex vivo* (Fig. 4, G to I). Whole-mount staining in synovial tissue explants confirmed expression of NGFR, TRKA, TRKB, and TRKC in mural cells within synovial microvasculature (Fig. 4J). Interestingly, stimulation of synovial tissue explants with NGF, BDNF, or NT3 resulted in markedly increased expression of a-smooth muscle actin (aSMA) in mural cells compared with vehicle-treated controls, indicating expansion of mural cell populations (Fig. 4K; Fig. S9A). Together, our data revealed a novel function of neurotrophins in regulating fibroblasts to mural cell differentiation. Whereas NGF preferentially promotes fibroblasts to pericyte differentiation, BDNF and NT-3 preferentially promotes fibroblasts to VSMC differentiation.

### NOTCH3 sensitize fibroblasts to NGF through transactivation of NGFR

To investigate the sequence of neurotrophin signaling during fibroblast to mural cell differentiation, we first analyzed NGF expression in fibroblasts and found that NGF expression colocalized with NOTCH3 in fibroblast-endothelial co-cultures (Fig. 5A-B), suggest that NOTCH3 initiates neurotrophin signaling by inducing NGF expression in fibroblasts adjacent to endothelial cells. Consistent with the hypothesis that NGF can be induced by NOTCH signaling, fibroblasts stimulated with DLL4 upregulated NGF expression (1.2-fold, *p* = 0.01) (Fig. 5C and Fig. S8A). Pharmacologic inhibition of NOTCH signaling via y-secretase inhibitor DAPT, or genetic deletion of *NOTCH3* by CRISPR-Cas9, abolished DLL4-induced *NGF* expression (Fig. 5, D and E). These data indicate that NOTCH3 is a critical initiator of neurotrophin signaling in fibroblasts through induction of NGF.

**Figure 5.**
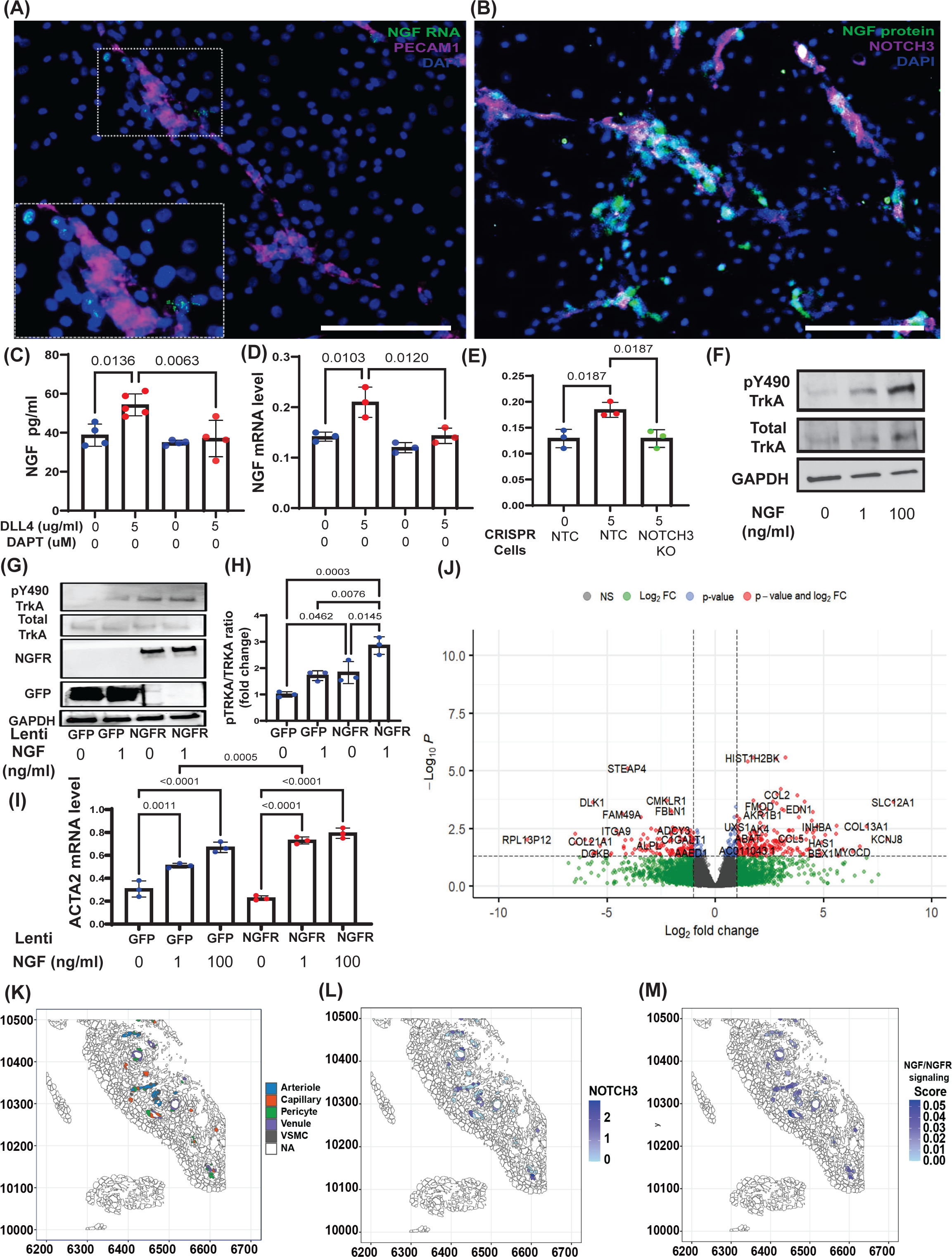
NOTCH3 regulates NGF-TRKA signaling in fibroblasts. Human fibroblasts were co-cultured with human umbilical vein endothelial cell line (HUVECs). Endothelial cells (HUVECs) were identified by PECAM1 expression (magenta). (A) RNAscope images showing transcript expression of NGF (green) and endothelial cell marker PECAM1 (magenta), overlaid with DAPI (blue). (B) Immunocytochemistry images showing NGF (green) and NOTCH3 protein (green), overlaid with DAPI (blue). Scale bars, 150 µm. All images were acquired at 20x magnification and cropped to the same scale. (C-D) Fibroblasts were treated with or without the y-secretase inhibitor to block NOTCH signaling, with or without DLL4, and NGF expression was measured. (C) NGF secretion (pg/ml). (D) NGF mRNA expression.(E) Fibroblasts with NOTCH3 knockout (NOTCH3 KO) generated using CRISPR-Cas9 were treated with or without DLL4, and NGF expression was measured. Individual data points represent biological replicates, with P values indicated. Statistical analysis was performed using a two-tailed Student’s t test for comparisons between two groups and one-way ANOVA for comparisons among multiple groups. Post hoc pairwise comparisons were conducted using t tests with Bonferroni correction to control for multiple comparisons. (F) Fibroblasts were treated with NGF (1 ng/ml or 100 ng/ml) or left untreated for 2 hours, and TRKA phosphorylation was assessed. Immunoblots showing phosphorylated TRKA (pY-TRKA), total TRKA, and GAPDH (loading control). (G-H) NGFR was overexpressed using a lentiviral CMV-driven system, with GFP as a control. NGFR- or GFPoverexpressing fibroblasts were cultured for 48 hours and treated for 1 hour with NGF (1 ng/ml or 100 ng/ml). (G) Immunoblots showing pY-TRKA, total TRKA, NGFR, GFP, and GAPDH. (H) Quantification of TRKA phosphorylation. (I) QRTPCR analysis of mural cell marker ACTA2 in GFP and NGFR overexpressed fibroblasts stimulated 1 and 100 ng/ml NGF treatment. (J) Bulk RNA sequencing analysis of differentially expressed genes in NGFR-overexpressing versus GFP-control fibroblasts treated without NGF (100 ng/ml). (K-M) NGFR-related gene signature enrichment was quantified in RA synovial tissue using Xenium spatial transcriptomics and the UCell rank-based scoring method. (K) Spatial mapping of vascular cell populations in RA synovial biopsy tissue. (L) Spatial expression of NOTCH3. (M) Spatial distribution of the NGF/TRKA/NGFR signaling score across RA synovial tissue

Because NTRK1/TRKA expression in synovial tissue and in cultured fibroblasts is extremely low (Fig. 2C and Fig. 3C), we next asked how fibroblasts can sense NGF upon its induction by NOTCH signaling. Because it has been suggested that the binding affinity of NGF to TRKA is potentiated through a NGFR-TRKA complex(33), we hypothesized that NGFR expression in fibroblasts could similarly potentiate TRKA signaling upon binding to NGF, providing a mechanisms by which fibroblasts can be sensitized to NGF despite expressing low level of *NTRK1*/TRKA. Consistently, NGF stimulation at a low concentration of 1 ng/ml resulted in minimal TRKA Y490 phosphorylation compared to maximal TRKA Y490 phosphorylation with high concentration of 100 ng/ml (Fig. 5F; Fig. S8B-C). In contrast, fibroblasts stably overexpressing NGFR (Fig. S8, D and E) exhibited a marked increase in TRKA Y490 phosphorylation at baseline and with 1 ng/ml of NGF stimulation (1.25-fold, *p=0.04*) (Fig 5, G and H). Consistent with enhanced sensitivity to NGF, NGFR-overexpressing fibroblasts exhibit elevated expression of pericyte-specific markers *ABCC9* (2.2-fold, *p=0.04*) and *RGS5* (1.2-fold) in response to 1 ng/ml of NGF, a level of induction comparable to the response observed with 100 ng/ml NGF in control GFP-transduced fibroblasts (Fig. S8F and G). NGFR-overexpressing fibroblasts also exhibited marked increased sensitivity to NGF-induced *ACTA2* expression (Fig. 5I). RNA-sequencing of NGFR-overexpressing fibroblasts revealed upregulation of multiple mural cell-associated genes, including *KCNJ8*, *ABCC9*(*26*), and *MYOCO* (*27*) in response to NGF stimulation (Fig. 5J). To determine whether the gene expression program driven by NGF/NGFR is present in RA synovial microvasculature, we generated an NGF/NGFR gene signature score based on 461 upregulated genes from RNA-sequencing and quantified its enrichment in RA synovial tissue using by projecting a cell-wise rank-based scoring approach (UCell) to RA synovial tissue transcriptomic data (Fig. 5K to M). We consistently observed enrichment of NGF/NGFR gene score in pericytes and VSMCs near capillaries and arterioles (Fig. 5L). Notably, NGF/NGFR gene score were co-localized with expression of NOTCH3 (Fig. 5M), supporting a role for NOTCH3 in initiating neurotrophin signaling in RA microvasculature. Together, these findings identify NOTCH3 as the key initiator of neurotrophin signaling by inducing NGF production in fibroblasts and sensitizing TRKA signaling through *NGFR* transactivation.

### TRK inhibitors reverse abnormal vascular maturation in RA synovial explants

Our data so far implicates neurotrophins as a novel regulator of RA vascular maturation through their ability to induce neighboring fibroblast to differentiate into mural cells. Importantly, TRK inhibitors are a class of FDA-approved targeted therapy in oncology for patients with NTRK gene fusions(*28*)(*29*). Our findings therefore raises the possibility of repurposing FDA-approved TRK inhibitors for RA therapy by targeting abnormal synovial vascular maturation. To evaluate this possibility, we first confirmed NOTCH signaling is required to initiate and sustain the activity of neurotrophin pathway in RA synovial microvasculature. In RA synovial explants, inhibition of NOTCH signaling by DAPT resulted in depletion of mural cells, as evidenced by reduction of aSMA expression (0.65-fold, *p* = 0.021) (Fig. 6A). Spatial transcriptomic analysis further revealed reduced vascular density, marked by decrease in *PECAM1* expression, in DAPT-treated RA synovial explants. Importantly, *NGF* expression was also reduced in RA synovial explants treated with DAPT (Fig. 6B), supporting the hypothesis that NOTCH signaling is required to initiate and sustain neurotrophin signaling in RA.

**Figure 6.**
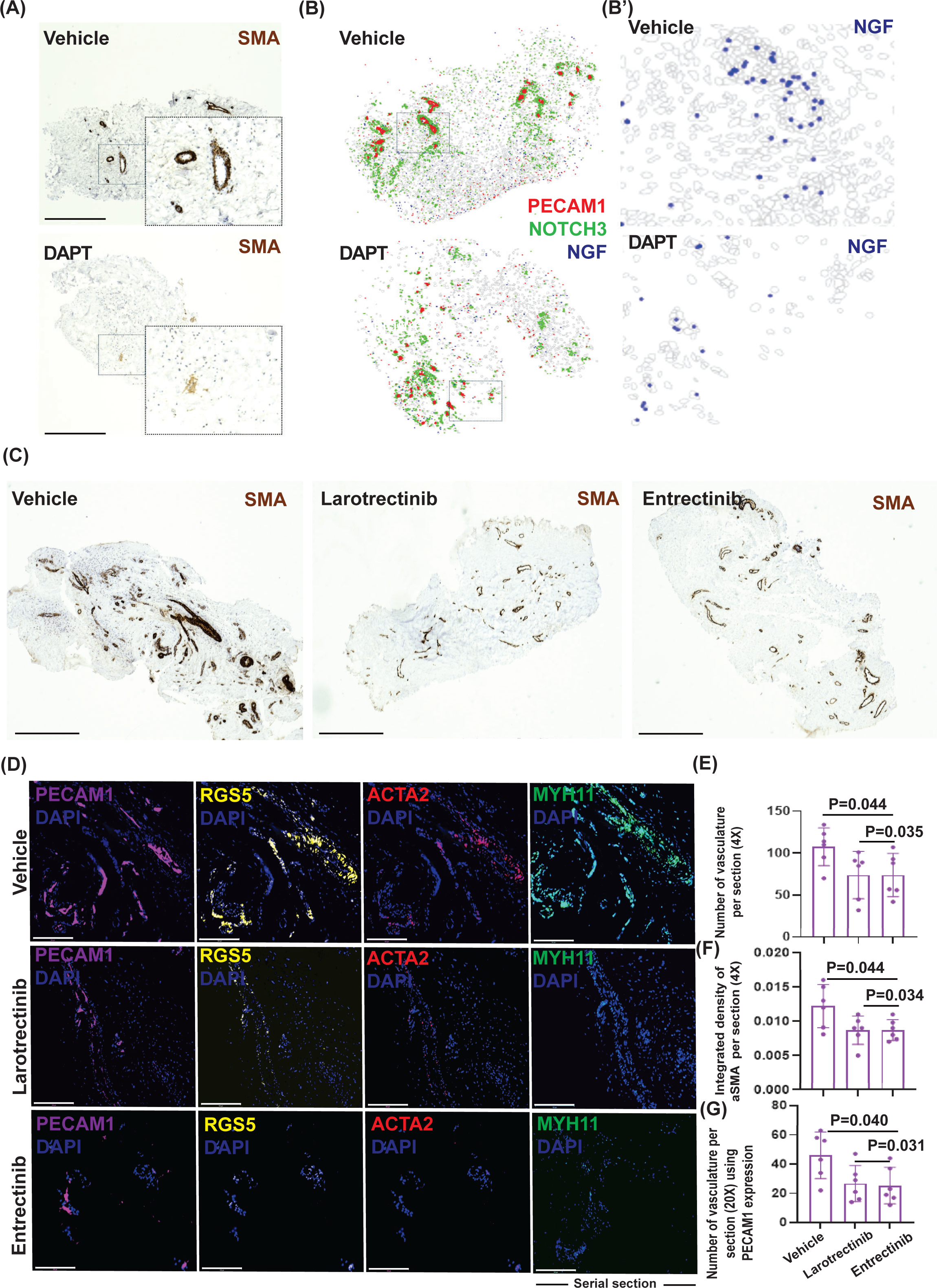
Reversal of pathological vascular maturation in RA synovial explants by TRK inhibitors. (A) Synovial tissue explant were embedded in Matrigel and treated with the NOTCH inhibitor DAPT. Organoids were fixed and stained for a-smooth muscle actin (aSMA) to visualize mural cells. (B) Spatial transcriptomic (Xenium) analysis of synovial explants treated with DAPT, showing expression of PECAM1 (red), NOTCH3 (green), and (B’) NGF (blue). (C) Representative images of synovial explants treated with entrectinib or larotrectinib, showing aSMA staining to visualize mural cells. Scale bar, 650 µm. (D) RNAscope images showing expression of RGS5 (yellow), ACTA2 (red), and MYH11 (green), overlaid with DAPI (blue), following entrectinib or larotrectinib treatment. For MYH11 staining either serial section or different sections were used. (E) Quantification of aSMA-positive vascular structures per tissue section following entrectinib or larotrectinib treatment. (F) Quantification of integrated aSMA staining density in synovial explants treated with entrectinib or larotrectinib.(G) Quantification of PECAM1-positive vascular structures per tissue section from 20x images following entrectinib or larotrectinib treatment. Scale bar, 150 µm. All images were acquired at 20x magnification and cropped to the same scale. Values are presented as mean ± standard deviation (SD). Individual data points represent biological replicates. Statistical analysis was performed using a two-tailed Student’s t test for comparisons between two groups and one-way ANOVA for comparisons among multiple groups. Post hoc pairwise comparisons were conducted using t tests with Bonferroni correction to control for multiple comparisons.

We next asked if the maturity of RA microvasculature can be modulated directly by agonists or antagonists of TRKs. Treatment with a TRKB agonist 7,8-dihydroxyflavone increased aSMA expression (1.3-fold, *p* = 0.021), whereas a pan-neurotrophin receptor agonist LM22B-10 produced the most robust increase (1.46-fold, *p* = 0.016) (Fig. S9C and D). In contrast, inhibition of TRKA or TRKB signaling via selective small-molecule inhibitors GW441756 and ANA-12, respectively, reduced aSMA expression (TRKAi, 0.7-fold, *p* = 0.18; TRKBi, 0.7-fold, *p* = 0.04) (Fig. S9 E and F). The strongest suppression of vascular mural cell fate was observed following treatment with a pan-TRK inhibitor GNF5837 (0.65-fold, *p* = 0.04) or DAPT (Fig S9F; Fig. 6A). Finally, we tested two FDA-approved TRK inhibitors, larotrectinib and entrectinib (*30, 31*), for their ability to reverse abnormal vascular maturation in RA. In cultured fibroblasts, neither drugs exhibited evidence of cytotoxicity at concentrations below 100 µM (Fig. S9B). Remarkably, both larotrectinib and entrectinib significantly reduced vascular density and aSMA expression in RA synovial explants (Fig. 6C). Quantitative analysis demonstrated a reduction in vascular area of 27% (*p* = 0.004) with larotrectinib and 24% (*p* = 0.035) with entrectinib. aSMA intensity was reduced by 36% (*p* = 0.044) and 40% (*p* = 0.034), respectively (Fig. 6 E). RNAscope analysis further confirmed reduced expression of mural cell markers following administration of TRK inhibitors, including *MYH11, RGS5*, and *ACTA2* (Fig. 6D). Quantification of vascular density using PECAM1 revealed a marked reduction following treatment with larotrectinib (54%, *p* = 0.04) and entrectinib (50%, *p* = 0.031)(Fig. 6 E). Together, these findings demonstrate that neurotrophin signaling sustains vascular maturation in RA synovia, and that TRK inhibitors effectively reverses vascular maturation and reduces pathological vascualrization in RA.

## Discussion

Pathological synovial vascularization plays a crucial role in sustaining chronic inflammation in RA(*32*), (*33*). Recent studies have highlighted novel roles of synovial vasculature in mediating various aspects of RA pathology, including inducing pathogenic fibroblast differentiation(*14*) and facilitating sensory nerve growth(*15*). In particular, synovial capillaries have been implicated as a driver of synovial tissue fibrosis which leads to poor treatment outcome in RA(*1*). Here, we provide, for the first time, a comprehensive, spatial transcriptomic analysis of synovial microvasculature from pre- and post- treatment to uncover a novel role of neurotrophin signaling in sustaining pathological vascular remodeling in RA. Our approach underscore the persistence of mature vascularization in RA despite treatment with conventional csDMARD or TNFi therapy. Mural cells play a crucial role in vascular maturation, stabilization, and function. Pericytes and VSMCs are distinct yet closely related mural cell types that associate with specific endothelial subtypes, with pericytes supporting capillary and venular endothelium and VSMCs stabilizing arterial vessels (*34*) (*35*). Neurotrophins are most known for their crucial functions during neural development(*19*). Genetic studies in animals have suggested a role for neurotrophins in vascular biology, as TRKB-null mice(*36*) exhibit defects in pericyte migration and VSMC function, while NT3-null mice(*37, 38*) display vascular abnormalities and impaired cardiac morphogenesis. Consistent with these observations, our studies utilizing pharmacological TRK agonists induce mural cell differentiation *ex vivo*, whereas TRK inhibitors reverse this effect. Our data indicate that neurotrophin signaling is tightly controlled by NOTCH3 receptor signaling (Fig. 7). Because NOTCH3 amplifies NOTCH signaling through a positive feedback loop via transcriptional induction of the NOTCH ligand JAGGED1(*39*), coupling of NOTCH3 and neurotrophin signaling enables mural cells to sense a dynamic range of endothelial NOTCH ligand availability, resulting in distinct neurotrophin signaling outputs. This coupling provides a mechanism by which endothelial cells dictate mural cell differentiation during maturation of venous and capillary endothelial cells, which express lower levels of NOTCH ligands, preferentially induce a pericyte phenotype through NGF-NGFR/TRKA signaling. Whereas higher levels of NOTCH ligands during maturation of arterioles promote VSMC differentiation phenotype via BDNF-TRKB and NT3-TRKC signaling.

**Figure 7:**
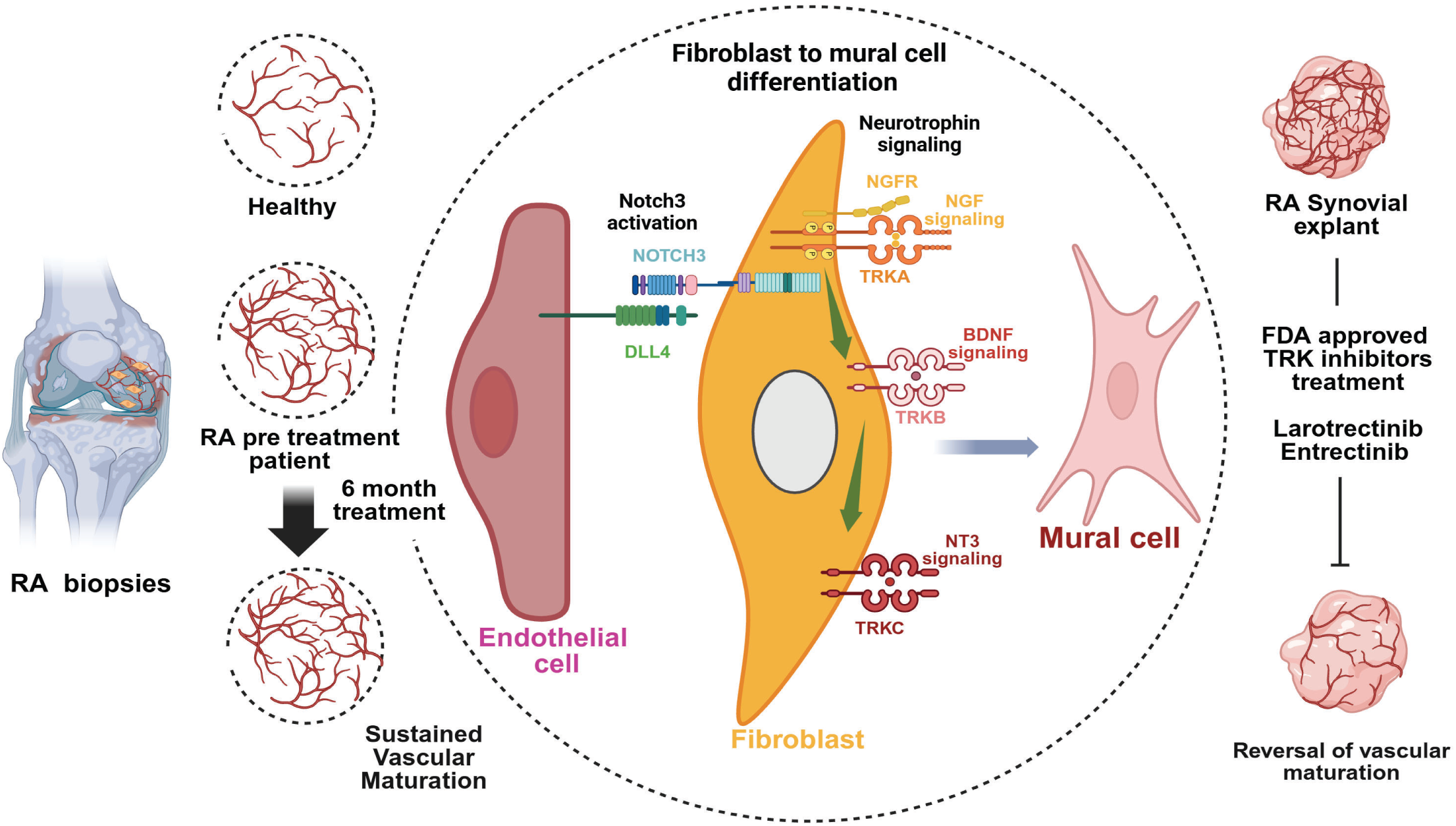
Schematic representation of a model illustrating NOTCH3-driven neurotrophin regulation of mural cell differentiation from fibroblast in pathological synovial vascular maturation. Pathological synovial vascular maturation in rheumatoid arthritis (RA) is actively sustained despite immunosuppressive therapy and persists independently of clinical remission. Spatial transcriptomic analyses reveal continued expansion of capillary, arteriolar, and mural cell populations, identifying the vasculature as a treatment-resistant disease compartment. Mechanistically, endothelial cells direct mural cell specification through NOTCH3-dependent induction of fibroblast-derived neurotrophin signaling. Pharmacologic inhibition of NOTCH or TRK signaling reduces mural cell activation and vascular density, with FDA-approved TRK inhibitors larotrectinib and entrectinib markedly attenuating pathological vascular maturation in human RA synovial explants. Together, this model identifies a NOTCH3-neurotrophin axis that sustains pathological synovial vascular maturation and represents a therapeutic target in treatment refractory RA.

Our finding that stromal cells are a major source of neurotrophic factors in RA synovia is consistent with the recent study illustrating the role of synovial fibroblasts in mediating sensory nerve growth and pain in RA(*15*). Indeed, our data supports the hypothesis that synovial fibroblasts -derived neurotrophin signaling may induce the coupling of blood vessels and sensory nerves in RA synovia. Nerve growth factor (NGF), in particular, is a well-established mediator of sensory nerve induction and sensitization, and its production by synovial fibroblasts suggests a direct mechanism by which fibroblast may promote neuronal growth in RA. Synovial fibroblast-derived neurotrophins may therefore coordinate vascular remodeling with neural infiltration, contributing to structural changes and pain sensitization. Our findings suggest that fibroblast-derived neurotrophin signaling may simultaneously sustain pathological vascular remodeling and may contribute in amplifying inflammatory signaling and pain perception in refractory RA. Further studies are needed to define the role of fibroblast-derived neurotrophins in recruiting sensory nerve growth in RA synovia, and whether or not the fibroblast neurotrophin signaling and the coupling of vascularization and innervation contributes to disease activity burden in refractory RA.

Current RA therapies target immune cells, inflammatory cytokines, or activation pathways downstream of cytokine receptors(*40*). Our study demonstrates that abnormal vascular maturation persists despite immunosuppressive therapy. These findings suggest that persistent microvascular remodeling in RA as a distinct pathological nidus that is not being effectively targeted by immunosuppressive therapies. One limitation of our study is that the treatment duration between the pre- and post-treatment biopsy is 6 months, therefore it remains possible that longer immunosuppressive treatment duration may result in normalization of synovial microvasculature in RA. Despite intense ongoing efforts in identifying a therapeutic strategy targeting stromal cells (*41*), no FDA-approved therapy directly targeting stromal cells exists. In the present study, pharmacologic modulation of neurotrophin signaling was sufficient to alter mural cell fate and vascular structure in established human RA microvasculature. Notably,

FDA-approved TRK inhibitors larotrectinib(*42*) and entrectinib(*43*), reduced vascular density in RA synovial explants, highlighting an opportunity to repurpose FDA-approved TRK inhibitors for RA patients.

### Materials and Methods Human Research

Human synovial tissue samples were obtained from Brigham and Women’s Hospital (MGB IRB no. 2019P002924) and Flinders Medical Center (Protocol#396.10)(*1*). Synovial tissue collected after patients undergoing arthroplasty or synovectomy procedures were dissected into 1-5 mm tissue fragments. These fragments were fixed in 10% formalin overnight at 4C, then stored in 70% ethanol at 4C before embedding in paraffin to make FEPE blocks for sectioning(*14*). For some tissue samples tissue fragments were submerged in Cryostor CS10 (#CS 10, Sigma-Aldrich), and stored at -80°C before long-term storage in liquid nitrogen. These tissues were used for either organoid studies or to make synovial fibroblast cell lines.

### Xenium slide preparation

Xenium slides were prepared from FEPE blocks and processed following the manufacturer’s protocol (CG000760 Rev A, 10X Genomics). Probe hybridization was conducted according to the “Xenium Prime In Situ Gene Expression with optional Cell Segmentation Staining For FF and FFPE samples on Xenium Slides User Guide” protocol (CG000760, 10X Genomics). Probes from the Xenium Prime 5K Human Pan Tissue & Pathways Panel (PN-1000671, 10X Genomics) and custom add-on panels were used for hybridization(*1*). Following the hybridization step, slides were washed and incubated with a ligation reaction mix for DNA amplification(*1*).

For visualizing cell boundary, cell segmentation staining was done using cell stain segmentation. Then the slides were stained with DAPI to visualize nuclei and treated with an autofluorescence quencher to minimize background interference. The prepared slides were then loaded onto the Xenium instrument, adhering to the guidelines outlined in the “Xenium Analyzer User Guide” (CG000584 Rev F, 10X Genomics). After scanning, the slides underwent post-run H&E staining as per the “Xenium In Situ Gene Expression - Post-Xenium Analyzer H&E Staining” protocol (CG000613 Rev A, 10X Genomics).

### Xenium Data Cell Typing and Niche Analysis

Initial single-cell typing analysis of Xenium data was performed as outlined in our earlier published work(*1*) and expanded the cohort to 22 patients (each with two time point samples) and included 2 healthy donors. After segmenting the cells (*44*), single-cell analysis was performed using Seurat v5.0.0 for quality control and data analysis. We thresholded the high quality cells based on transcripts and features per cell, and kept 2 million high quality cells across the 46 samples, and then we robustly typed lineages and fine vascular cell states with our integrative annotation procedure as follows: we combined both Xenium and AMP scRNA-seq using the 5,063 overlapped genes, and normalized the data by log-transforming the transcript counts for each sample, with the median number of detected transcripts from both Xenium and AMP scRNA-seq reference serving as the scaling factor, computed the top 50 PCs of the normalized feature by cell expression matrix, and integrated over sample-specific effects in the PC using Harmony v1.2.4 (*1*). We then estimated a weighted nearest neighbor graph of the cells in harmonized PC space and used the graph to estimate a two-dimensional UMAP embedding. We used the same graph to partition the cells by maximum modularity with the Louvain algorithm [cite Louvain] with resolution 0.3. We next identified cluster markers with differential expression analysis. To do this, we used the Wilcoxon rank sum test as implemented in the presto package(*45*). Cell type labels were assigned to each cluster based on known function state and lineage markers. At the lineage level, we classified cells into lining fibroblast, sublining fibroblast, mural, endothelial, adipocyte, myeloid, T/NK, B/plasma cells. For downstream analysis, endothelial cells, and mural cells were subset from the cells, and fine subtype based on the same procedures above. This resulted in 368,217 vascular cells across 46 samples include 130,108 pericytes, 97,717 venule, 56,699 capillary, 40,359 arteriole cell, 41,827 VSMC cell, and 1507 endothelial lymphatic cell.

An analogous cell-type clustering analysis was performed on the 50-gene and 377-gene custom panel datasets derived from cultured explant vascular tissue. Both datasets were processed using the same analytical pipeline. The resulting spatial dataset comprised 114,058 cells obtained from a single rheumatoid arthritis (RA) synovial biopsy.

### Single cell signature scoring

We performed bulk RNA-seq for the organoid with drug treatment and without drug treatment, and identified the drug response markers with the differential expression analysis; the top differentially expressed gene sets were used to define NGFR-related gene markers. Both the markers and normalized Xenium gene expression were provided to compute the NGFR enrichment scores using a R package UCell (https://github.com/carmonalab/UCell, UCell and pyUCell: single-cell gene signature scoring for R and python), which utilized marker expression rank-based scoring statistics that robustly measure the signature score distribution in cell types.

### Cell culture

Synovial fibroblast cell lines were generated from the synovial tissue mentioned above using an established protocol.(*14*) Briefly, after rinsing the tissue in PBS, it was cut into small pieces and collected in a 50 mL Falcon tube using sterile forceps. The tissue pieces were then incubated in a digestion enzyme solution containing Dispase II (Sigma, #494207801) at 100 µg/mL, DNase I (Sigma, #10104159001) at 100 µg/mL, and Liberase TL (Sigma, #5401020001) at 100 µg/mL in plain RPMI for 1 hour on a shaker at 37°C. Following digestion, 30 mL of complete FLS medium (DMEM supplemented with 10% fetal bovine serum, HEPES, MEM amino acids, L-glutamine, penicillin-streptomycin, nonessential MEM amino acids, 2-mercaptoethanol, and gentamicin) was added to inactivate the enzymes. The cell suspension was then passed through a 70 µm filter and centrifuged at 1500 rpm for 10 minutes. Cell pellets were resuspended in complete FLS medium, and the medium was changed every 3 days. Synovial fibroblast cell lines were cultured and maintained (3 to 6 passages for experiments) in complete FLS media as described above HUVECs (Thermofisher) were cultured and maintained in EGM2 media consisting of EGM-Plus media (Lonza # CC-5035) supplemented with the EGM-plus bulletkit (Lonza,cc-3162).

### Cell Culture Studies

Synovial fibroblasts were cultured in DMEM supplemented with 10% fetal bovine serum (FBS), as previously described(*46*). For Notch activation experiments, fibroblasts were seeded onto culture plates coated with recombinant DLL4-Fc (10185-D4, R&D Systems) overnight at 4°C at a concentration of 5 µg/ml (or the concentration indicated for each experiment). After coating, fibroblasts were seeded on DLL4-coated or vehicle-coated plates and cultured for 24-72 hours. For neurotrophin (NT) stimulation, recombinant NGF (256-GF, R&D Systems), BDNF (11166-BD, R&D Systems), and NT3 (267-N3-005, R&D Systems) were reconstituted in DMSO or PBS and diluted in media. Fibroblasts were treated with NTs at the following concentrations: NGF (1, 100 ng/ml), BDNF (100 ng/ml), and NT3 (50, 100 ng/ml). Small-molecule inhibitors, including GW-441756 (#2238, Tocris, TrkA inhibitor), ANA12 (#4781, Tocris, TrkB inhibitor), GNF 5837 (#4559, Tocris TrkA/BC inhibitor), Entrectinib (Cat. No.: HY-12678, MedChemExpress) Larotrectinib (Cat. No.: HY-12866, MedChemExpress) were used at concentrations of 1, 5, and 10 µM respectively. Cell viability checked using WST-1 (Cat No. 501594400, Milliore sigma). Neurotrophin agonists LM22B 10 (#6037, Tocris, TrkB/C agonist) and 7-8 DHF (#3826, Tocris, TrkB agonist) were used at 1, 5, and 10 µM. respectively. For co-cultures, plates were pre-coated with a 1:10 dilution of matrigel (Corning, Cat. 356231) in PBS.Then HUVECs ( passage 3-7) were cultured with synovial fibroblasts at 1:1, 1:5, or 1:10 ratios in EGM2 media for 72-96 hours, in the presence or absence of neurotrophins, neurotrophin modulators(list them or say as described above), and the Notch inhibitor DAPT (#2634, Tocris, 10 µM), as indicated above. For siRNA experiments, fibroblasts were transfected with siRNAs targeting Notch3 or neurotrophin receptors (NTRK1, NTRK2, NTRK3) using RNAiMax reagent (Life Technologies), as previously described(*46*). Cells were transfected for 2 days with various silencer select siRNA (NGFR assay ID-S194655: NTRK1 assay ID-S534734: NTRK2 assay ID-n321595: NTRK3 assay ID-s9753, NOTCH3-106100), following manufacturer protocol (thermo scientific), and subsequently treated with various neurotrophins. NOTCH3 KO cells were generated using P3 primary cell 4D-nucleofector X kit S (#V4XP-3032, Lonza), according to the manufacturer’s protocol. Guide RNAs were designed using Synthego design tool(*1*).

### Synovial Tissue explant and micromass studies

Synovial tissue-derived organoids were generated from synovial tissue collected as described above. Synovial tissue was thawed and washed in PBS and DMEM media, and 2mm punch biopsy was used to extract cylindrical sections of synovium. The synovial biopsies were embedded into 96-well plates pre-coated with 50 µL of ice-cold Matrigel (Corning #356231) and incubated at 37°C for 30 minutes to allow gel solidification. After solidification, organoids were cultured in EGM2 (Lonza #CC-5035) media and after 2 hours of incubation, were treated with various neurotrophins and their modulators. Following 3 days of treatment, organoids were fixed in formalin for immunohistochemistry (IHC) and RNAScope experiments. Histological analysis of synovial organoids was performed by fixing organoids with 4% paraformaldehyde, embedding in paraffin, and staining with hematoxylin/eosin (Sigma, H&E), as previously described(*47*),(*^14^*). For Fibroblast micromass 200,000 synovial fibroblasts were mixed with 35 µL droplet of Matrigel (Corning #356231) and cultured on polyHEMA (#25249-16-5, Sigma-Aldrich)-coated plates in EGM2 media (as described above) for 3-7 days. Micromasses were treated with various neurotrophins, and images were acquired after 3 days. For fibroblast-endothelial cell organoids, 100,000 synovial fibroblasts and 100,000 HUVECs were combined in a single 35 µL Matrigel droplets and cultured on polyHEMA (Sigma-Aldrich)-coated plates in EGM2 media for 14-21 days. Cells were labeled with fluorescent dyes PKH67(Red, #PKH67GL, Sigma-Aldrich) and PKH26 (Red, #PKH26GL, Sigma-Aldrich) to visualize fibroblast and HUVEC cells. Micromasses were treated with various neurotrophins and images were acquired after 3 days.

### Whole-mount immunofluorescence staining

Synovial organoids were collected after treatment and washed with PBS to remove excess matrigel. Organoids were fixed in 4% formalin at RT for 20 minutes. After washing, organoids were blocked with 1% BSA in a permeabilizing solution (0.1% Triton-X) for 1-2 hours at RT. After washing, organoids were incubated with primary antibodies NGFR, NTRK1, NTRK2, NTRK3 (Cell Signaling Technology, #4638), MYH11 (Proteintech, #21404-1-AP), and CD31(Biolegend, #303106) overnight at 4°C. The following day, organoids were rinsed with PBS and incubated with secondary antibodies (Alexa fluor antibodies: AF555 anti-rabbit, #A-21424, AF488 anti-rabbit, #A11034, anti-goat #A110055, and AF647 anti-rabbit, #A32733). After secondary antibody incubation, organoids were washed and stained with DAPI to visualize nuclei. Images were acquired on an EVOS M7000 at 1.25x and 4X magnification and analyzed using ImageJ software.

### RNAscope analysis and quantification

Synovial tissue sections were processed using the manufacturer’s protocol with an RNAScope multiplex fluorescent V2 assay (ACD Bio, SOP 45-009A). Different probes for neurotrophins and mural cell markers were used (provided in Table 2) and Images were acquired on an EVOS M7000. Similarly for co-culture studies, fibroblasts and endothelial cells were seeded on matrigel as described earlier. Fibroblast cells cultured for 2 days and transfected with siRNA for 48 hours. Then fibroblast cells were cultured with endothelial cells. After culture and treatment with siRNA, cells were washed with PBS, fixed in 10% NBF, and processed f according to the manufacturer’s protocol with an RNAScope multiplex fluorescent V2 assay (ACD Bio, SOP 45-009A). DAPI is used to sustain nuclei. Images were acquired on an EVOS M7000. Results were quantified by segmenting nuclei based on DAPI staining using Cellpose(*48*) and using scikit-image(*49*) to perform nuclear expansion to approximate cell boundaries and calculate mean intensities per cell. Per cell intensities of PECAM1, RGS5, and MYH11 were normalized by estimated cell area. Cell types were assigned by gating normalized per cell intensity on high (>= 90th percentile) and low (<90th percentile) marker expression. Cells with high PECAM1 expression were labeled as endothelial cells, cells with high RGS5 and low MYH11 as RGS5+, low RGS5 and high MYH11 as MYH11+, RGS5 and MYH11 high as MYH11+/RGS5+, and RGS5 and MYH11 low as MYH11-/RGS5-. Distance to the nearest endothelial cell was calculated with RANN(*50*).

### Western Blot

Fibroblasts were cultured and treated with various neurotrophins (NGF-1, 100ng/ml: BDNF-100ng/ml, NT3-100ng/ml). Cell lysates were prepared using a RIPA buffer supplemented with protease and phosphatase inhibitor mini-tablets (Thermo Fisher Scientific, #A32961) and a phosphatase inhibitor cocktail (Active Motif, #37492). For phosphorylation studies, lysates were collected after 2 hours of treatment. For functional analysis, lysates were collected after 72 hours of treatment. Protein concentration was determined using the Pierce BCA Protein Assay Kit (Thermo Fisher Scientific, #23227), and 30-50 µg of protein was subjected to SDS-PAGE on mini-protein TGX precast Gels (Bio-Rad#45610954) and transferred to PVDF membranes (Bio-Rad#1620174). Using the Bio-Rad TRANS-BLOT TURBO transfer system (Bio-Rad, #690BR324) with a dry transfer protocol for 30 minutes according to the manufacturer’s instructions (Bio-Rad). Following transfer, membranes were blocked for 20 minutes in Everyblot blocking buffer (Bio-Rad # 12010020) and incubated overnight at 4°C with primary antibodies from TrkA and TrkB antibody sampler kit against TrkA, TrkB, p-TrkA/TrkB(Cell Signaling Technology, #4638, 1:500), CNN1 (Proteintech-24855-1-AP), MYH11 (Proteintech-21404-1-AP), NGF (Abcam-ab52918) and GAPDH (Thermo Fisher Scientific, #MA5-15738),or beta-actin (Cell Signaling Technology, #3700). Following primary antibody incubation, membranes were incubated with HRP-conjugated secondary antibodies (Thermo FisherScientific, anti-Rabbit #32460, anti-Mouse #31430, or anti-Goat#A16005) for 1 hour at room temperature. All the antibody dilutions were in Everyblot blocking buffer. Blots were developed using SuperSignal™ West Femto Maximum Sensitivity Substrate (Thermo Fisher Scientific,#34095) or SuperSignal™ West Pico PLUS Chemiluminescent Substrate (Thermo Fisher Scientific, #34577) and imaged using a Bio-Rad ChemiDoc imaging system.

### Immunohistochemistry and Imaging Analysis

Immunohistochemistry (IHC) was performed on formalin-fixed, paraffin-embedded (FFPE) synovial tissue sections. Tissue sections were processed as described earlier.(*14*) Primary antibodies against smooth muscle actin (SMA), PECAM, NGFR, TRKA, TRKB, and TRKC (as described above) were used according to standard protocols at Brigham and Women’s Hospital Pathology Core. Nuclei were stained with DAPI, and slides were mounted using VectaMount® Permanent Mounting Medium (Vector Laboratories. #H-5000-60). Images were acquired using the EVOS M7000imaging system. Images were analyzed using imageJ software.

### Lentiviral Protocol

To generate lentiviruses, 6 x 10⁶ 293FT cells were seeded in a T75 culture flask containing 10 mL of complete FLS media (DMEM supplemented with 10% fetal bovine serum, HEPES, MEM amino acids, L-glutamine, and nonessential MEM amino acids, without antibiotics). Cells were incubated overnight at 37°C in a humidified 5% CO2incubator. The next day, the lentiviral vector mix was prepared according to the Virapower™ HiPerform™Lentiviral FastTiter™ Gateway® Expression protocol (Thermo Fisher Scientific, #K534000).Briefly, 1.95 mL of Opti-MEM and 55 µL of Lipofectamine™ 3000 reagent were mixed and incubated at room temperature for 5 minutes. In a separate tube, 1.95 mL of Opti-MEM, 17.8 µg of packaging mix, 11.18 µg of lentiviral plasmid containing the NGFR gene with N-terminal GFP-tag(pLV-BsdCMV-hNGFR, VectorBuilder, #VB230823-1657hvq) or a control plasmid pLV-Bsd-CMV-EGFP (VectorBuilder, # VB230502-1039MVR) and 47 µL of P3000™ reagent were mixed and incubated for 5 minutes. Following these incubations, the DNA mix was added to the Lipofectamine™ 3000 mix, and the resulting mixture was incubated for an additional 20 minutes at room temperature. This DNA/lipofectamine mix was added to 293FT cells in DMEM with 5% FBS and no antibiotics. After 4 days, viruses were collected and concentrated using PEG-it ((System Biosciences, # LV-810-A-1). Synovial fibroblasts at the density of 1X10^5^ cells were plated in FLS media with no antibiotics for 2 days and then, transduced with 50 ul of viral particles and 10 ug/ml of polybrene. GFP expression in transduced cells was monitored microscopically.

### Real-time Quantitative PCR

Cells were treated and lysed in TRIzol (#15596026,Thermofisher) to extract RNA as per manufacturer protocols.cDNA was synthesized using QuantiTect Reverse Transcription Kit (#205311 Qiagen) according to manufacturer protocols. The qPCR mastermix was prepared using Brilliant III qRT-PCR Master Mixes (#5994-1166EN, Agilent) and samples were run on an AriaMX Real Time PCR machine (Agilent). mRNA levels were normalized to Glyceraldehyde-3-phosphate dehydrogenase (GAPDH) and calculated using the 2-ΔΔCT method(*51*). The primer list is provided in Table 1.

**Table 1:**
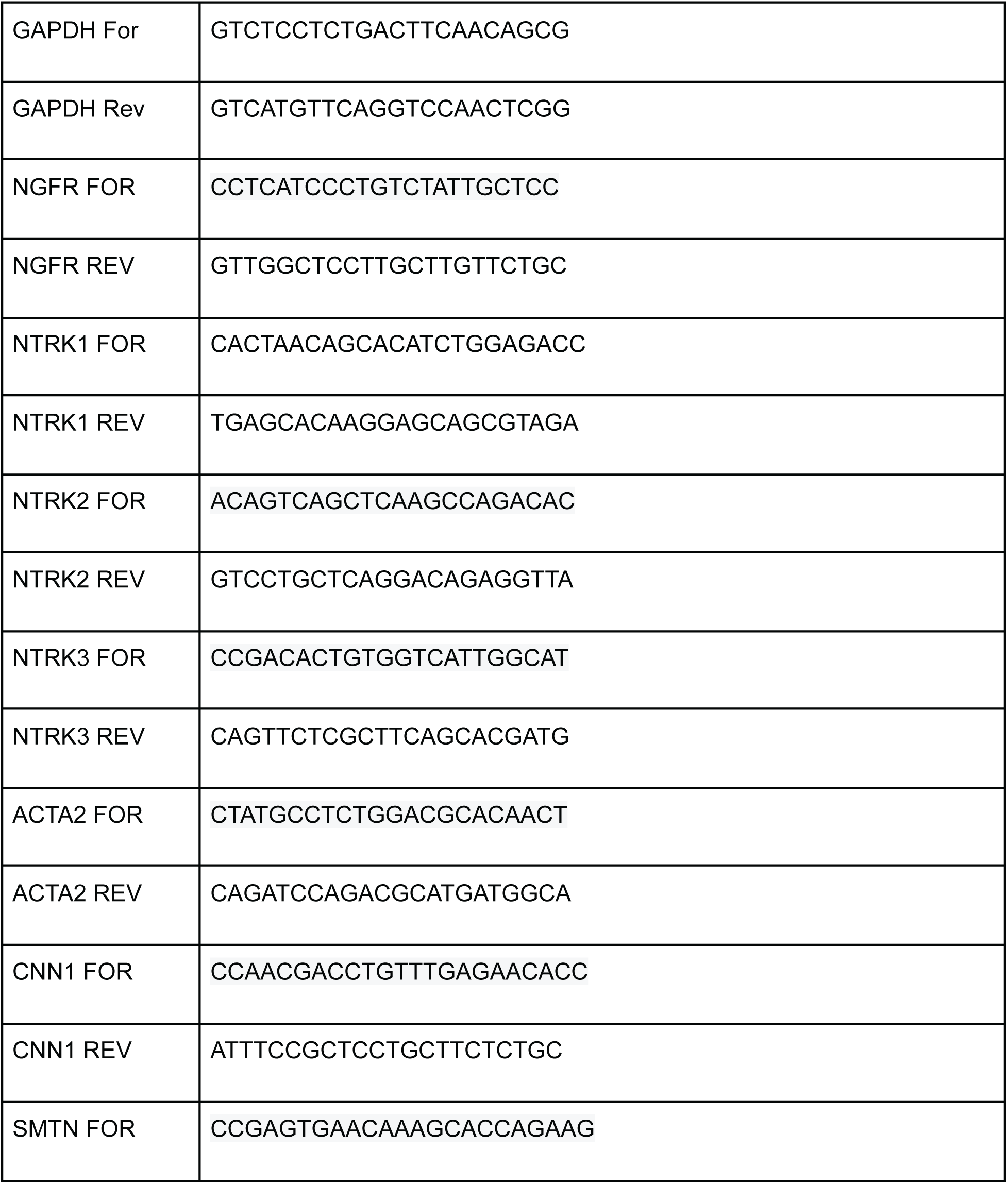
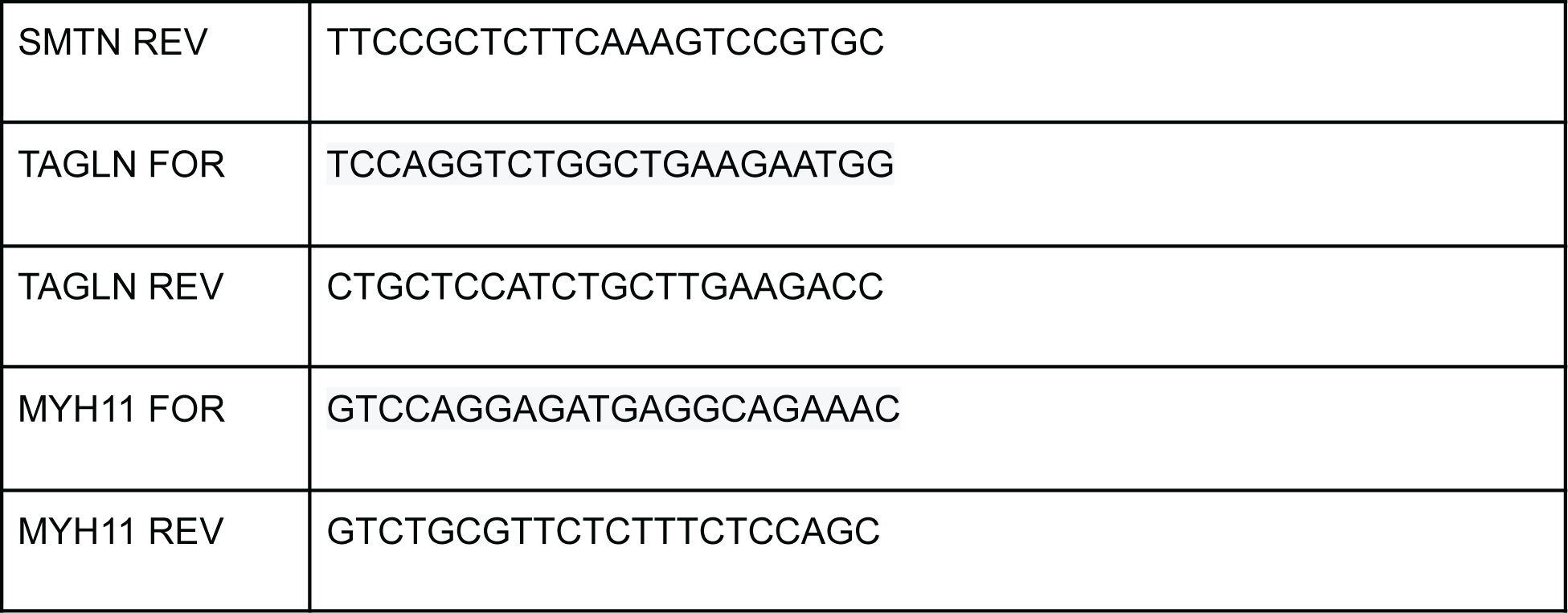
Human primer List.

**Table 2:** RNAScope probe List.

|  |  |
| --- | --- |
| Mouse MYH11 | 316101-Mm-Myh11 |
| Human NGF | 549451-C2-Hs-NGF-C2 |
| Human NGFR | 406331-Hs-NGFR |
| Human NTRK1 | 402631-C4- Hs-NTRK1-C4 |
| Human NTRK2 | 402621-C2-Hs-NTRK2-C2 |
| Human NTRK3 | 406341-C2-Hs-NTRK3-C2 |
| Human ACTA2 | 444771-C2-Hs-ACTA2-O1-C2 |
| Human MYH11 | 444151-C4- Hs-MYH11-C4 |
| Human NOTCH3 | 487381-Hs-PECAM1-O1-C3 |
| Human RGS5 | 402621-C2-Hs-RGS5 - C1 |

### Statistical Analysis

Statistical tests were performed using Graphpad Prism version 10.4.1. Statistical analyses were performed as indicated in the figure legends. All data were analyzed using appropriate tests for comparisons. P-values less than 0.05 were considered statistically significant. For statistical analysis, two-tailed student’s t-test was used. One-way ANOVA was conducted to compare the means across multiple groups. Following a significant result, post-hoc pairwise comparisons were performed using t-tests with Bonferroni correction to control for multiple comparisons.

### Software

Image analysis and quantification were performed using ImageJ. Single-cell analyses were conducted using R software. Graph generation and statistical analyses were performed using GraphPad Prism. Schematics were created with BioRender. Figures were assembled using Adobe software

---

**Fig. S1.**
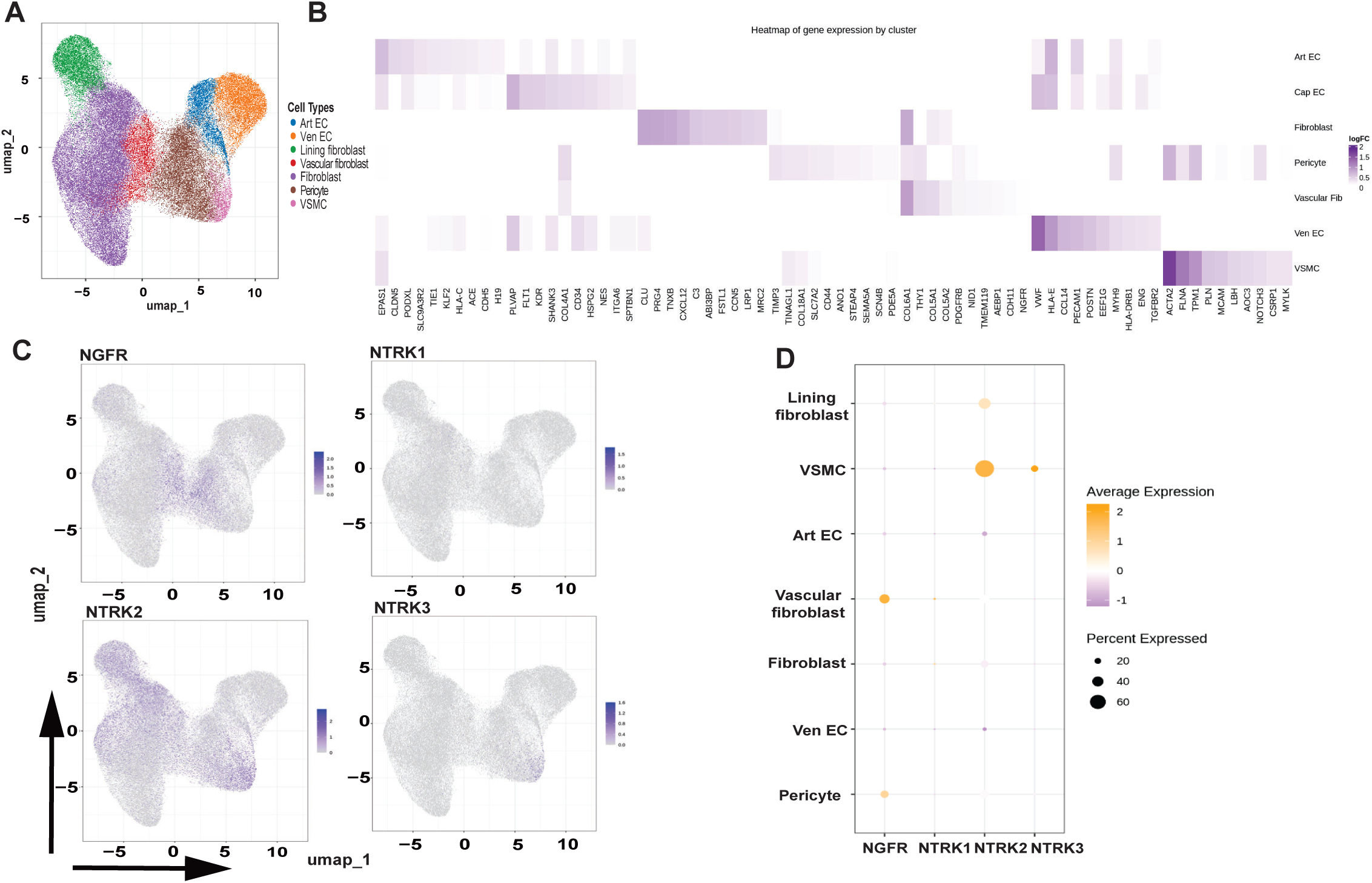
5ingle-cell RNA sequencing analysis of RA synovial biopsy samples using targeted 377- and 50-gene panels. (A) UMAP projection of all cells annotated with integrated vascular cell-type labels. (B) Heatmap showing expression of marker gene signatures used to define vascular cell clusters.(C) UMAP feature plots showing expression of neurotrophin receptors NGFR, NTRK1, NTRK2, and NTRK3 across vascular cell types.(D) Dot plot summarizing expression of neurotrophin receptors NGFR, NTRK1, NTRK2, and NTRK3 across annotated vascular cell populations.

**Fig. S2.**
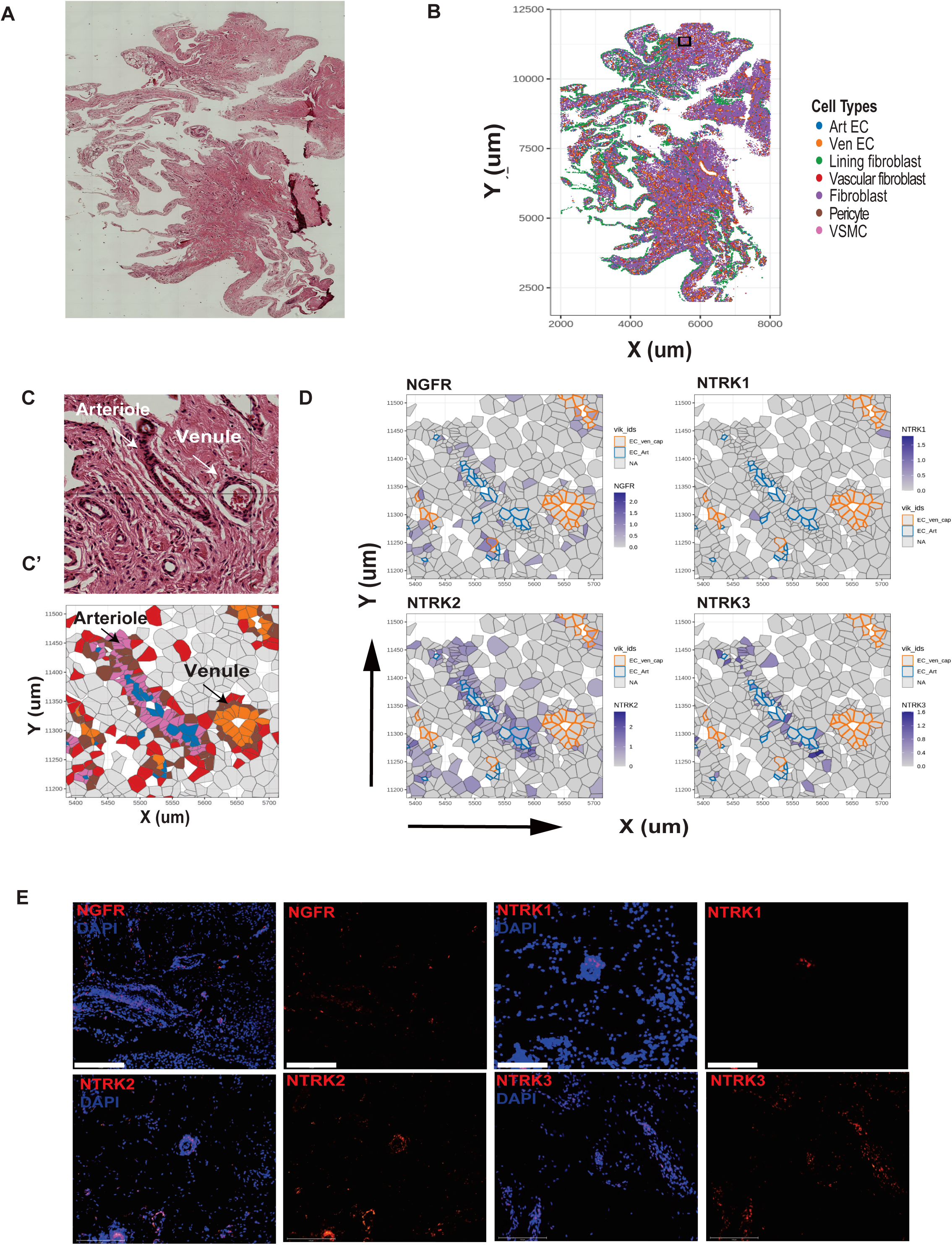
Spatial analysis and mapping of vascular cell populations in RA synovial biopsy samples using targeted 377- and 50-gene panels. (A) Histological hematoxylin and eosin (H&E) staining of RA synovial tissue biopsy.(B) Representative spatial transcriptomic plot of the synovial tissue biopsy colored by annotated vascular cell types. (C) H&E staining of a selected region of the synovial tissue biopsy. (C’) Corresponding high-resolution spatial transcriptomic plot of the same region colored by vascular cell types.(D) Spatial gene expression patterns of neurotrophin receptors NGFR, NTRK1, NTRK2, and NTRK3 across mapped vascular cell populations.(E) RNAscope images showing transcript expression of neurotrophin receptors NGFR, NTRK1, NTRK2, and NTRK3 (red) in RA synovial tissue sections, overlaid with DAPI (blue). Scale bar, 150 μm.

**Fig. S3.**
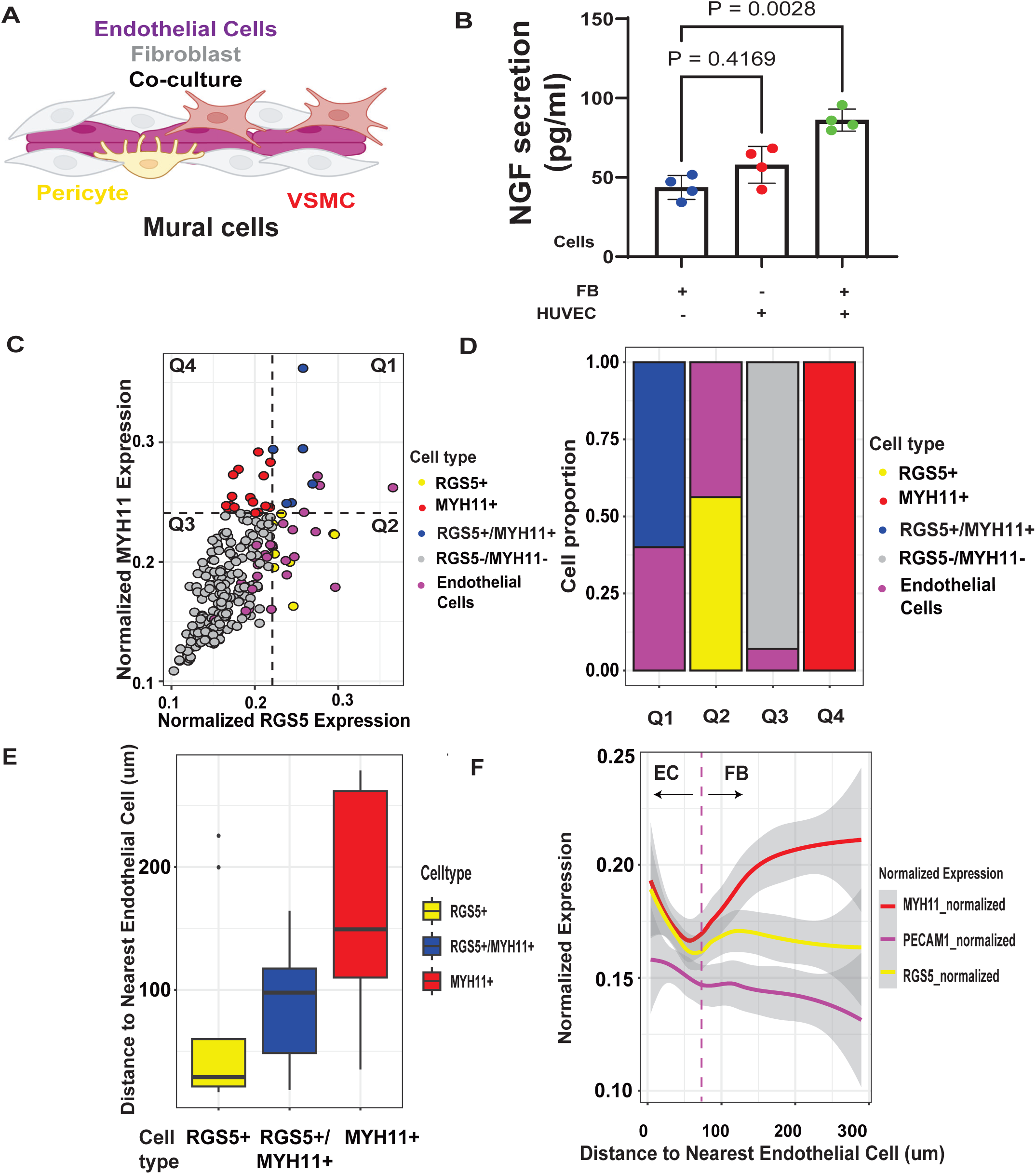
Endothelial–fibroblast co-culture induces spatially organized mural cell differentiation. (A) Schematic illustrating the endothelial cell–fibroblast–mural cell co-culture system.(B) ELISA quantification of NGF secretion levels (pg/ml) in co-culture.(C) Distribution of RGS5+, MYH11+, and RGS5+/MYH11+ double-positive cells.(D) Proportion of cells in each quadrant, with quadrant boundaries defined by RGS5 and MYH11 expression as described in Methods.(E) Distance of RGS5+ pericytes, MYH11+ VSMCs, and RGS5+/MYH11+ double positive cells from the nearest endothelial cells.(F) Normalized expression of RGS5, MYH11, and PECAM1 relative to distance from the nearest endothelial cells.

**Fig. S4.**
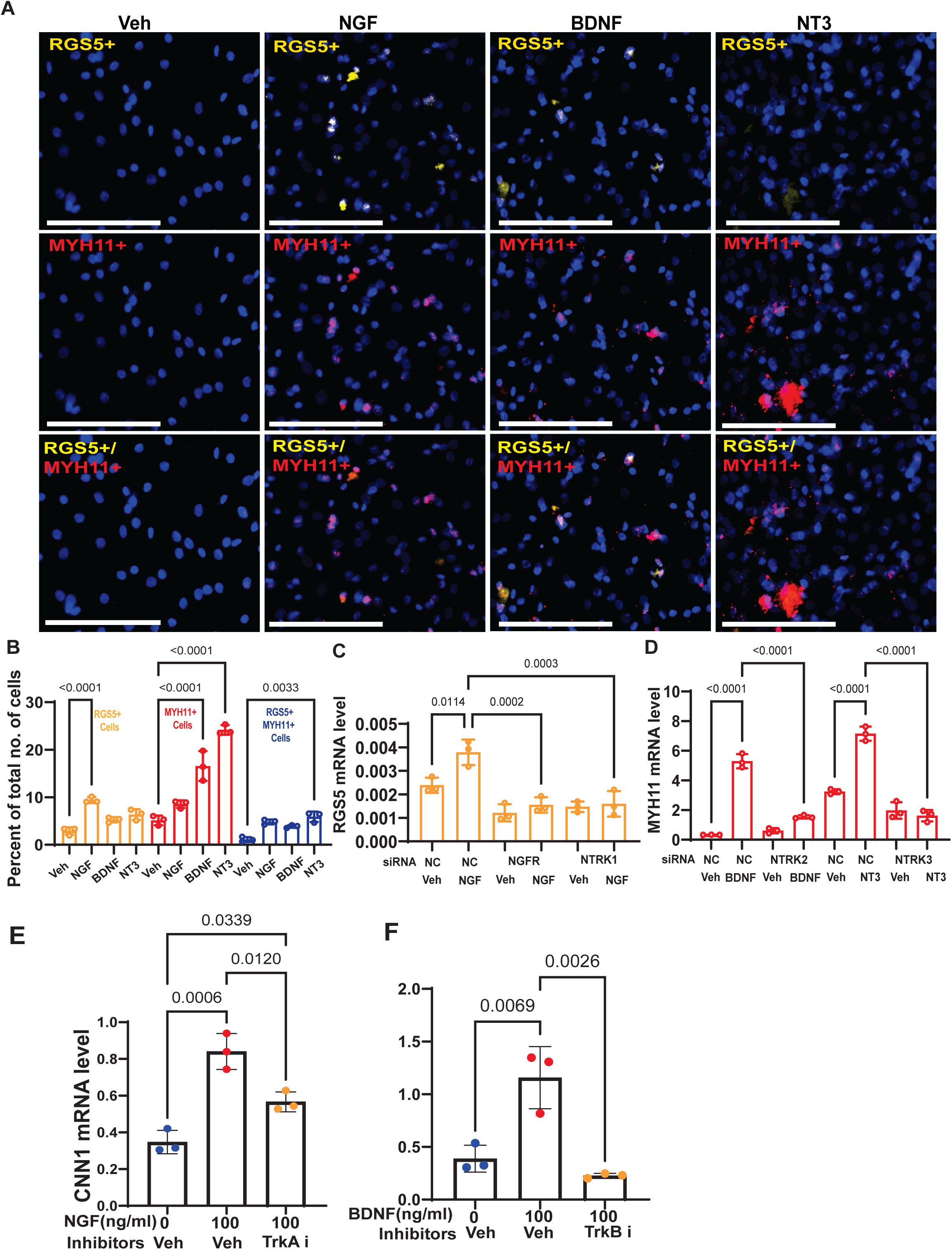
Neurotrophin signaling differentially regulates pericyte and VSMC marker expression. (A) RNAscope images showing transcript expression of pericyte marker RGS5 (yellow) and VSMC marker MYH11 (red) in fibroblasts treated with different neurotrophins, overlaid with DAPI (blue). Scale bar, 150 µm. All images were acquired at 20x magnification and cropped to the same scale.(B) Quantification of total numbers of RGS5+, MYH11+, and RGS5+/MYH11+ double-positive cells following treatment with the indicated neurotrophins.(C) mRNA expression of the pericyte marker RGS5 in fibroblasts transfected with siRNAs targeting NGFR or NTRK1, in the presence or absence of NGF stimulation.(D) mRNA expression of the VSMC marker MYH11 in fibroblasts transfected with siRNAs targeting NTRK2 or NTRK3, in the presence or absence of BDNF or NT3 stimulation, respectively.(E-F) mRNA expression of the mural cell marker CNN1 (calponin) in fibroblasts stimulated with NGF (E) in the presence or absence of a TRKA inhibitor, or stimulated with BDNF (F) in the presence or absence of an NTRK2 inhibitor.

**Fig. S5.**
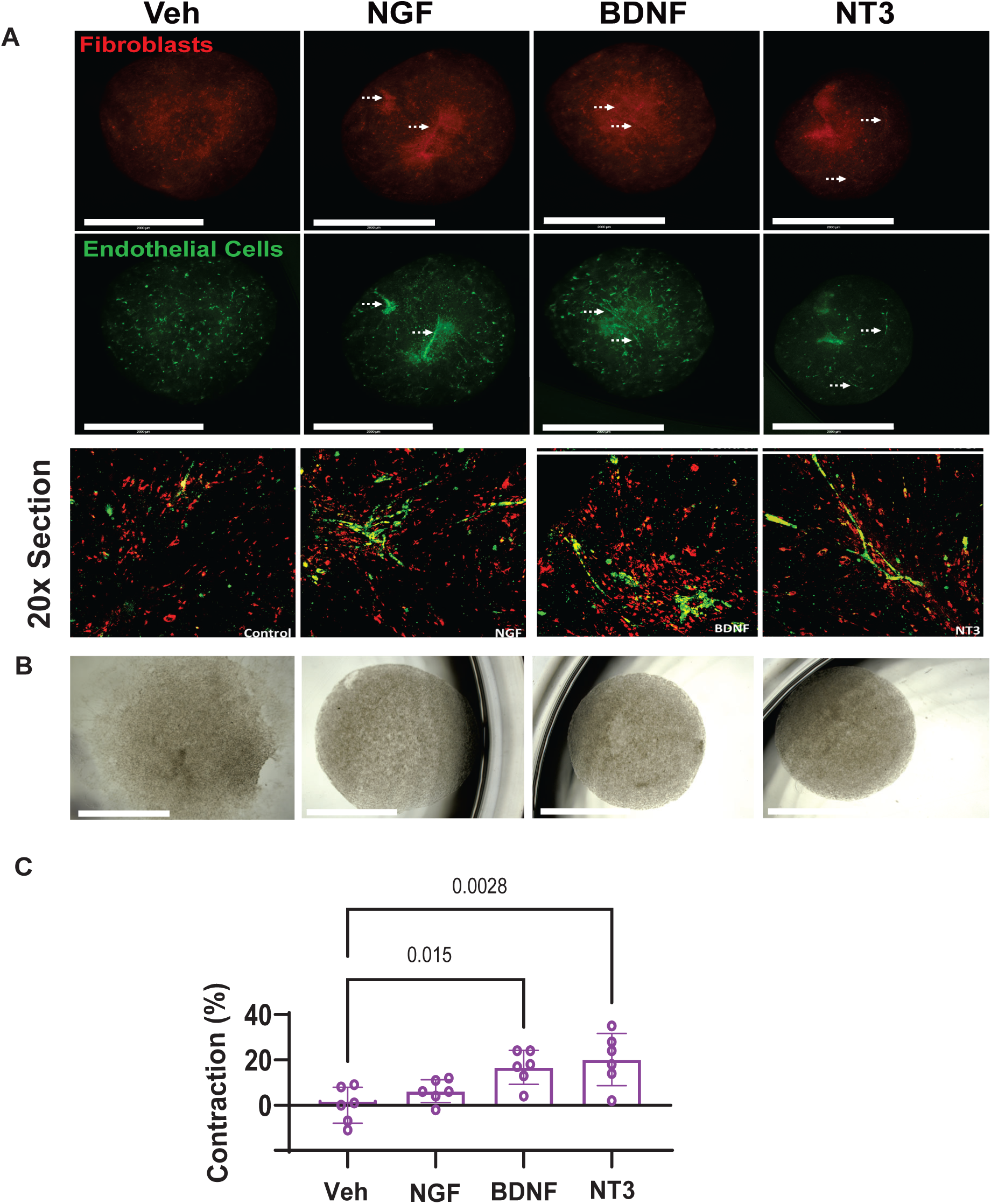
Neurotrophin stimulation promotes vascular tube-like structure formation and contractility in micromass organoids. (A) Micromass organoids generated in matrigel and cultured for 7 days. Fibroblasts (red) and endothelial cells (green) were stained to visualize cellular organization following treatment with the indicated neurotrophins. Scale bar, 2000 μm. Images were acquired at 1.25x magnification and cropped to the same scale. Lower panels show enlarged confocal (20X) images highlighting vascular- or tube-like structure formation.(B) Micromass organoids generated from fibroblast monocultures, shown as a control condition. Scale bar, 2000 µm.(C) Quantification of percent diameter contraction of micromass organoids. Values are presented as mean ± standard deviation (SD). Individual data points represent biological replicates. Statistical analysis was performed using a one-way ANOVA for comparisons among multiple groups. Post hoc pairwise comparisons were conducted using *t* tests with Bonferroni correction to control for multiple comparisons.

**Fig. S6.**
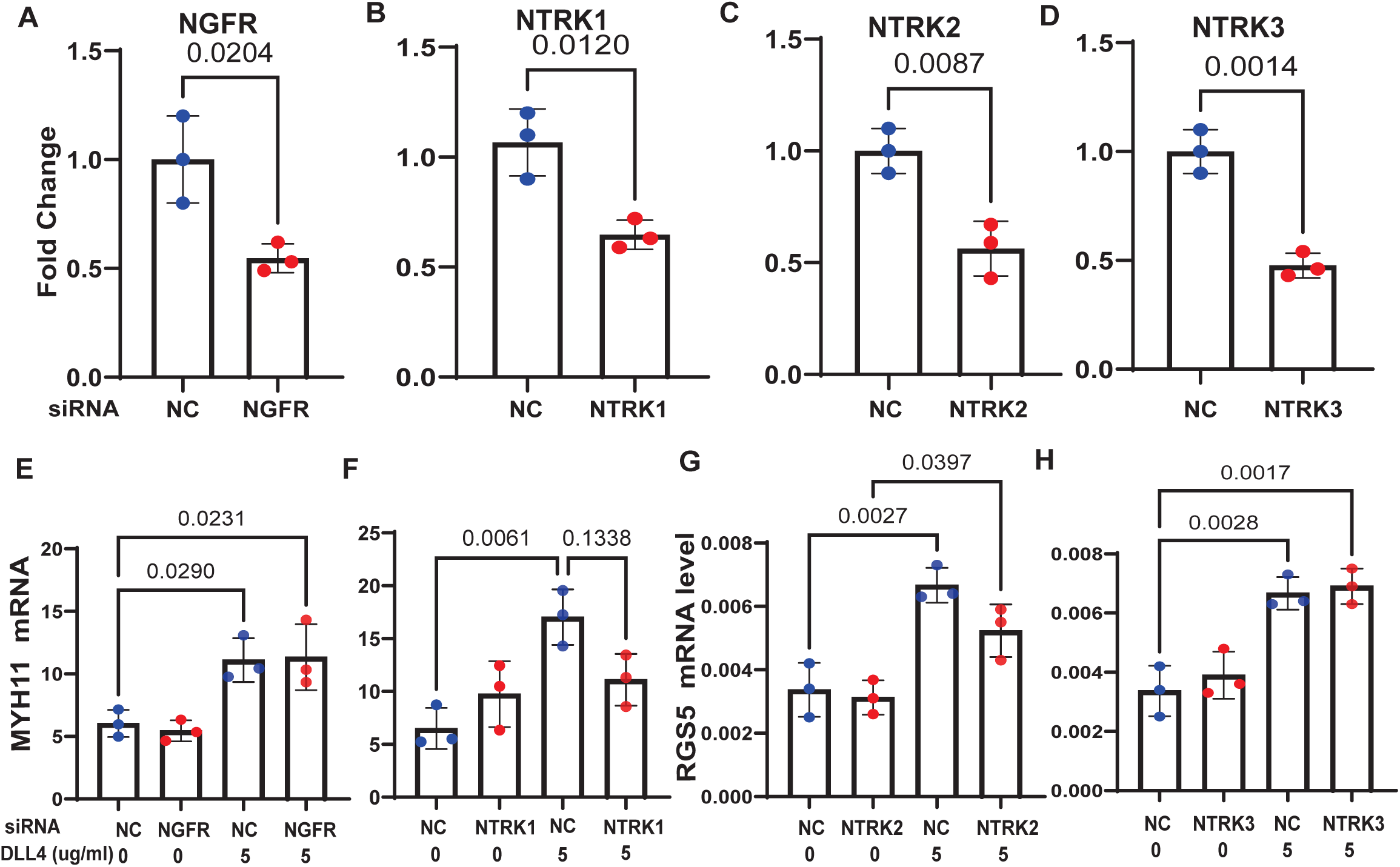
Validation of neurotrophin receptor knockdown and effects on mural cell marker expression. (A-D) mRNA expression levels of NGFR (A), NTRK1 (B), NTRK2 (C), and NTRK3 (D) in fibroblasts treated with siRNAs targeting the indicated genes, demonstrating knockdown efficiency. (E-H) Fibroblasts were treated with siRNAs targeting NGFR, NTRK1, NTRK2, or NTRK3 in the presence or absence of DLL4 (5 µg/mL), and expression of mural cell markers was assessed. MYH11 mRNA expression following knockdown of NGFR (E) or NTRK1 (F), and RGS5 mRNA expression following knockdown of NTRK2 (G) or NTRK3 (H).lndividual data points represent biological replicates, with P values indicated. Statistical analysis was performed using a two-tailed Student’s t test for comparisons between two groups and one-way ANOVA for comparisons among multiple groups. Post hoc pairwise comparisons were conducted using t tests with Bonferroni correction to control for multiple comparisons.

**Fig S7:**
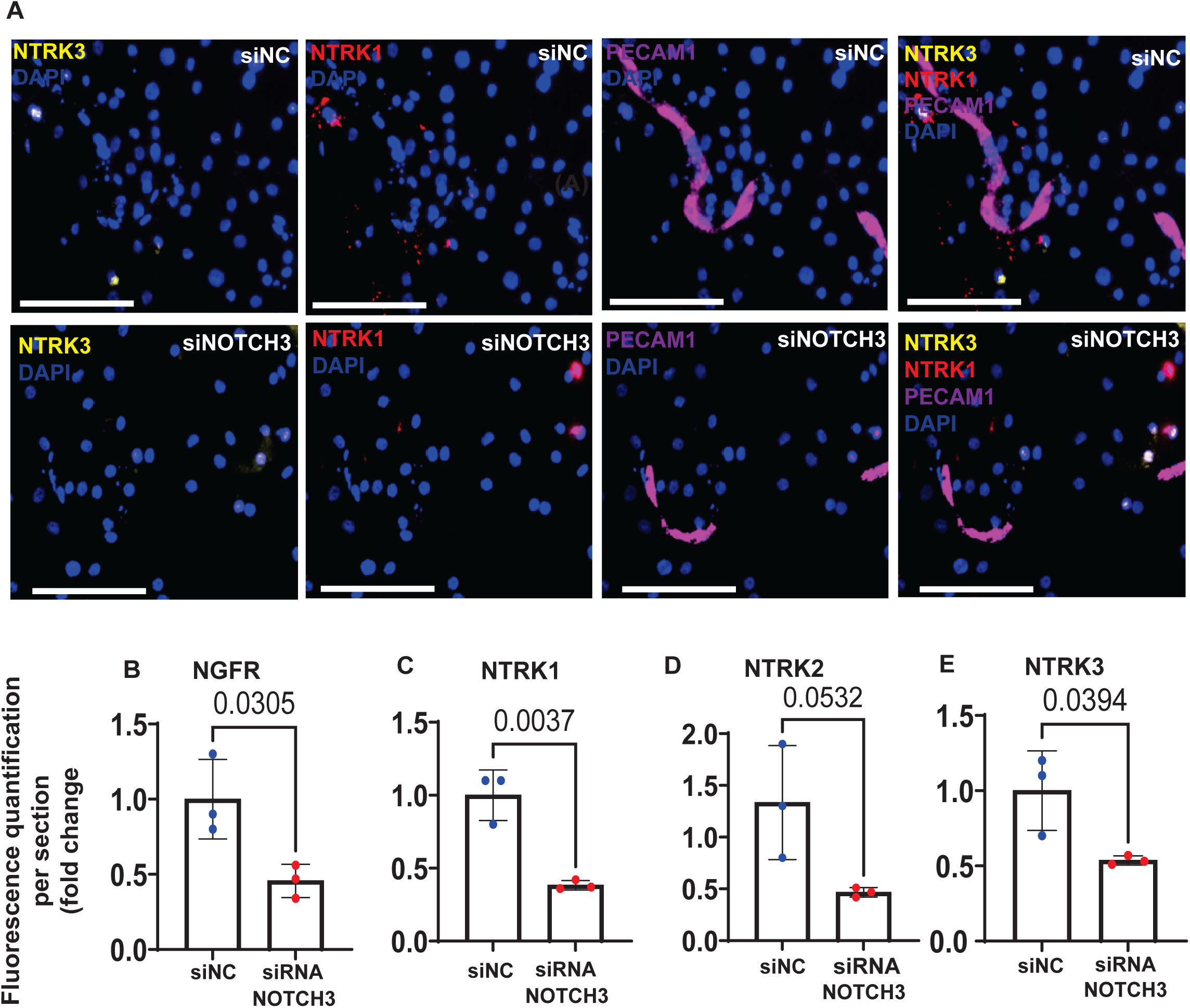
NOTCH3 regulates expression of neurotrophin receptor in fibroblasts. Fibroblasts were treated with NOTCH3 siRNA and co-cultured with endothelial cells. (A) RNAscope images show transcripts of NTRK1 (red) and NTRK3 (yellow) in the co-culture, overlaid on DAPI (blue). Scale bar, 150 μm. All images were acquired at 20× magnification and cropped to the same scale. (B-E) Fluorescence quantification per section of NGFR, NTRK1, NTRK2 and NTRK3. Individual data points represent biological replicates, with *P* values indicated. Statistical analysis was performed using a two-tailed Student’s *t* test for comparisons between two groups and one-way ANOVA for comparisons among multiple groups. Post hoc pairwise comparisons were conducted using t tests with Bonferroni correction to control for multiple comparisons.

**Fig. S8.**
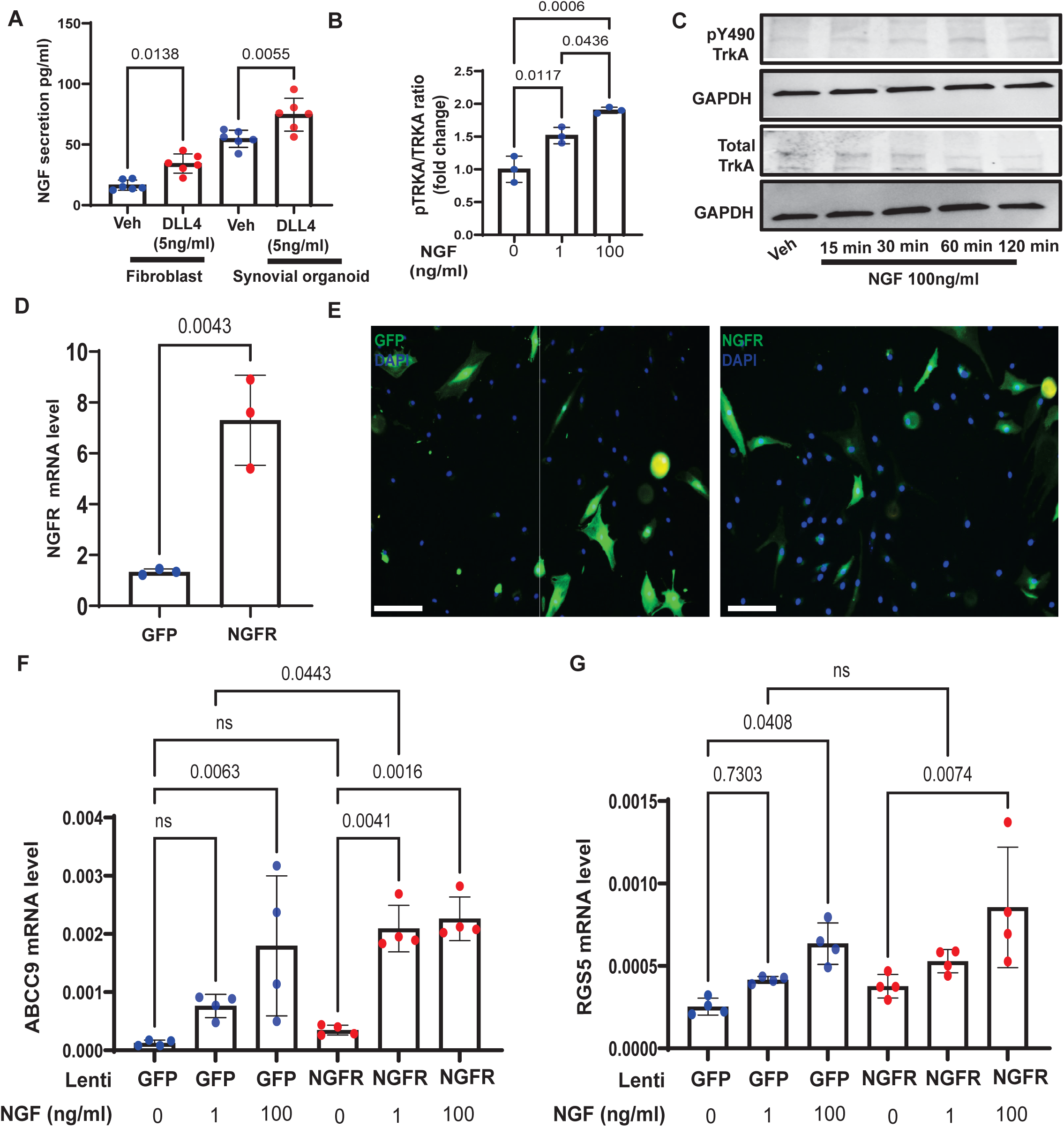
NOTCH and NGFR potentiate NGF–TRKA signaling in fibroblasts. (A) ELISA quantification of NGF secretion (pg/ml) in conditioned media collected from fibroblasts and synovial organoids treated with or without DLL4 (5 µg/ml).(B) Quantification of TRKA phosphorylation, shown as pY-TRKA relative to total TRKA. (C) Immunoblots showing pYTRKA, total TRKA, and GAPDH (loading control) in unstimulated fibroblasts or fibroblasts stimulated with NGF for the indicated time points (0, 15, 30, 60, and 120 min). (D-G) NGFR was overexpressed using a CMV-driven lentiviral system, with GFP used as a control. (D) mRNA expression levels of NGFR in fibroblasts. (E) Immunoluorescence images showing GFP expression in control cells and NGFR-overexpressing fibroblasts.(F–G) qRT-PCR analysis of pericyte-associated markers in GFP- or NGFR-overexpressing fibroblasts stimulated with NGF (1 ng/ml or 100 ng/ml). (F) ABCC9 expression. (G) RGS5 expression.Values are presented as mean ± standard deviation (SD). Individual data points represent biological replicates, with *P* values indicated. Statistical analysis was performed using one-way ANOVA for comparisons among multiple groups. Post hoc pairwise comparisons were conducted using *t* tests with Bonferroni correction to control for multiple comparisons.

**Fig. S9.**
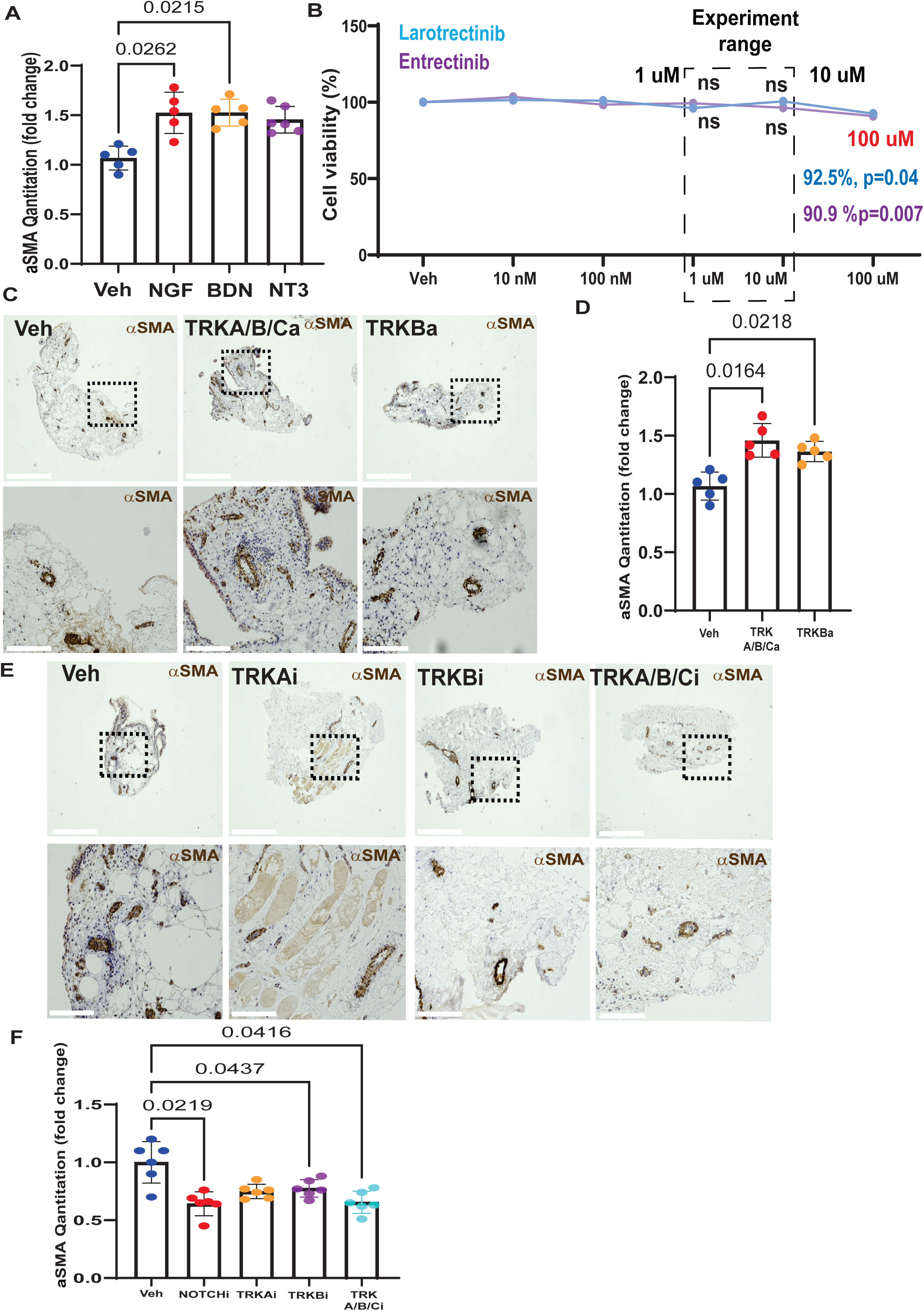
Pharmacologic modulation of neurotrophin signaling alters mural cell activation in synovial explants. (A) Quantification of a–smooth muscle actin (aSMA) staining in synovial organoids treated with neurotrophins (NGF, BDNF, NT3).(B) Cell viability assay of entrectinib and larotrectinib across a range of concentrations (0-100 uM). Values are presented as mean, with *P* values indicated (two-tailed student’s t-test). (C) Representative aSMA staining images of synovial organoids treated with neurotrophin receptor agonists. (D) Quantification of aSMA staining following treatment with neurotrophin receptor agonists (LM22B-10 and 7,8dihydroxylavone).(E) Representative aSMA staining images of synovial organoids treated with neurotrophin receptor antagonists. (F) Quantification of aSMA staining following treatment with neurotrophin receptor antagonists (DAPT, GNF5837, GW441756, and ANA-12). Values are presented as mean ± standard deviation (SD). Individual data points represent biological replicates, with *P* values indicated on the graphs. Statistical analysis was performed using one-way ANOVA for comparisons among multiple groups. Post hoc pairwise comparisons were conducted using *t* tests with Bonferroni correction to control multiple comparisons.

## Acknowledgment

We thank members of the Wei lab, Korsunsky lab, and Brenner lab for helpful discussions.This work is supported by a NIH NIAMS R01AR085028, Arthritis Foundation Rheumatoid Arthritis Research Program, a Burroughs Wellcome Fund Career Awards for Medical Scientists to K.W., a Brigham and Women’s Hospital Department of Medicine - Broad Institution collaborative research Award, a Brigham and Women’s Hospital Llura Gund Award for Rheumatoid Arthritis Research. K.W. is supported by an NIH-NIAMS K08AR077037.

## Author contributions

Conceptualization: V.K, and K.W. Experimental design and data generation: V.K. and S.K. Xenium data generation and panel design: G.C., S.A.P, V.K., K.B. Data analysis: Q.Q, M.T., A.B.R.M., V.K. Synovial tissue acquisition: S.P., V.K, P.E.B., J.K.L., M.H.J., M.D.W. Synovial tissue acquisition and relevant clinical and histological correlations: M.D.W. Figure generation: V.K, M.T., Q.Q. Writing original draft: V.K., M.T., and K.W. Draft reviewing and editing: all. Supervision: K.W. Funding acquisition: K.W.

